# Using Multivariate Pattern Analysis to Increase Effect Sizes for Event-Related Potential Analyses

**DOI:** 10.1101/2023.11.07.566051

**Authors:** Carlos Daniel Carrasco, Brett Bahle, Aaron Matthew Simmons, Steven J. Luck

## Abstract

Multivariate pattern analysis approaches can be applied to the topographic distribution of event-related potential (ERP) signals to ‘decode’ subtly different stimulus classes, such as different faces or different orientations. These approaches are extremely sensitive, and it seems possible that they could also be used to increase effect sizes and statistical power in traditional paradigms that ask whether an ERP component differs in amplitude across conditions. To assess this possibility, we leveraged the open-source ERP CORE dataset and compared the effect sizes resulting from conventional univariate analyses of mean amplitude with two multivariate pattern analysis approaches (support vector machine decoding and the cross-validated Mahalanobis distance, both of which are easy to compute using open-source software). We assessed these approaches across seven widely studied ERP components (N170, N400, N2pc, P3b, lateral readiness potential, error related negativity, and mismatch negativity). Across all components, we found that multivariate approaches yielded effect sizes that were as large or larger than the effect sizes produced by univariate approaches. These results indicate that researchers could obtain larger effect sizes, and therefore greater statistical power, by using multivariate analysis of topographic voltage patterns instead of traditional univariate analyses in many ERP studies.

## 1. Introduction

For decades, ERP researchers have assessed differences in amplitude between experimental conditions with univariate analyses. In these analyses, the mean or peak voltage within a time period is measured for each condition at one or more electrode sites, and univariate statistical analyses (e.g., *t* tests, analyses of variance) are used to determine whether the differences in amplitude across conditions or groups are statistically significant. Researchers often record from >20 electrodes, but the analyses typically focus on either one electrode or a cluster of electrodes where the component of interest is largest(Zhang & Kappenman, 2023). Researchers sometimes put separate measures from each electrode into a univariate statistical analysis and look for condition x electrode site interactions, but these types of analyses typically have low statistical power (Luck, 2014). Mass univariate analyses can also be used to identify electrode sites showing significant differences between conditions (Maris & Oostenveld, 2007), but they may also suffer from reduced power (Groppe et al., 2011). A major shortcoming of these univariate approaches in general is that they assume that all participants have approximately the same scalp distribution for a given component, which is typically not a good assumption given the large individual differences in cortical folding patterns (Borrell, 2018; Fernández et al., 2016; Kroenke & Bayly, 2018), which can produce substantial differences in scalp topography (Hajizadeh et al., 2021).

To address these shortcomings, researchers have begun to apply multivariate pattern analysis (MVPA) techniques to ERP data to gain insight into brain activity that may not be related to traditional ERPs (Ashton et al., 2022; Chan et al., 2011; Daly, 2023; de Vries et al., 2019; Grootswagers et al., 2017; Gurariy et al., 2022; Hubbard et al., 2019; D. Li et al., 2023; Murphy et al., 2011; Wang et al., 2012; Yang et al., 2021). In particular, machine learning algorithms can be used to distinguish slight differences between different stimulus classes in the pattern of voltage across electrode sites^1^. This makes it possible for a given algorithm to correctly guess the stimulus class that produced an ERP waveform from new data that were not used for training. In other words, the stimulus class is *encoded* by the brain and can be *decoded* by the algorithm. For example, MVPA approaches have been used to decode the identity and emotional expression of a face being perceived (Bae, 2021; Dobs et al., 2019; Nemrodov et al., 2018; Smith & Smith, 2019). Decoding can also be used to gain insight into a range of cognitive domains including motion perception, attention, and working memory (Bae & Luck, 2019; Bae, 2021; Bae & Luck, 2018; Ester & Weese, 2023; Y. Li et al., 2022; Nadra et al., 2023; Noah et al., 2020).

These approaches are extremely sensitive, capable of reliably distinguishing ERP patterns for different faces, different directions of motion, different orientations, and so on. It is therefore plausible that they could also be used to increase power for conventional ERP analyses.

The most common ERP decoding approach uses support vector machines (SVMs). As an example, we will briefly explain how this approach works for decoding whether a participant was shown a face stimulus or a car stimulus, and we will then describe how it can be used to answer traditional ERP questions (e.g., whether the N170 component is larger for faces than for cars). Figure 1a shows a simplified example of SVM decoding in which simulated single-trial data from only two electrode sites (e.g., PO8 and Cz) are used to decode whether the subject had been shown a picture of a face or a picture of a car. Each dot in the scatterplot shows the voltage from this pair of electrode sites for a given trial. Decoding is usually performed separately at each latency to take advantage of the ERP technique’s temporal resolution, so the data in Figure 1a would be from a specific latency (e.g., 170 ms after stimulus onset). Although faces and cars produce similar distributions of voltage at each electrode when considered individually, the SVM is able to find a *decision line* that can separate the face and car data. Note that this example greatly exaggerates the difference between conditions for the sake of clarity.

**Figure 1.**
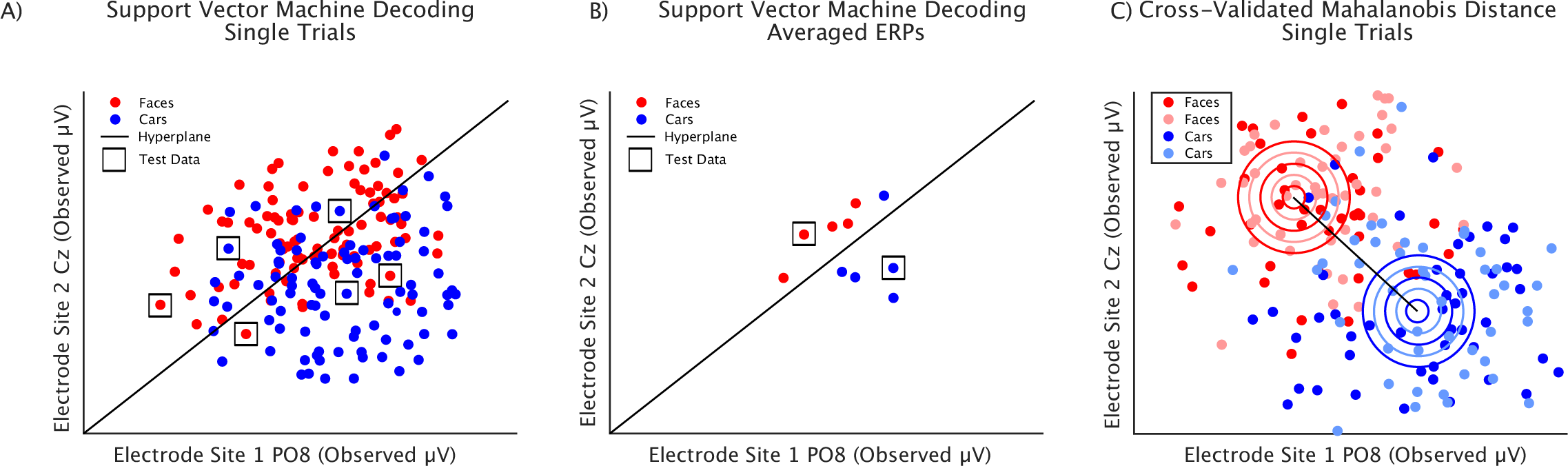
Simplified example of support vector machine (SVM) decoding and the cross-validated Mahalanobis distance (*crossnobis* distance). This example uses simulated data from only two electrode sites, which can be represented as a 2-dimensional space. The data are shown from a single latency. A) Single-trial decoding. Each dot represents the voltage at these two electrodes on a single trial. The SVM finds a decision line (*hyperplane*) that best divides the trials into the two classes. Once the SVM is trained, it is tested with data from trials that were not used to train the SVM (shown as dots surrounded by squares). The SVM classifies these test cases according to which side of the decision line they fall on. The proportion of these classifications that were correct (i.e., matched the true classes for these trials) is then used to quantify the accuracy of the decoder. B) Decoding from averaged ERPs. The 80 single trials for each stimulus class in the original dataset are divided into five sets of 16 trials and then averaged to create five averaged ERPs for each class. Four of the averaged ERPs from each class are used for training and the remaining averaged ERP from each class is used for testing. C) Crossnobis algorithm. This approach operates on single-trial data. It finds the centroid and multidimensional standard deviation for each class and computes the distance between the two means scaled by the standard deviations. The trials from each class are randomly divided into two subsets to allow for a cross-validation procedure. Here we are showing simulated single-trial data with centroids that are much more distant than would typically be observed in real data.

To avoid capitalizing on noise (i.e., *overfitting*), the SVM finds the decision line using a set of training cases and then is tested on cases that were not used for training (indicated by the squares in Figure 1a). A given test case is classified according to which side of the decision line it falls on, and this classification is then compared with the true class to determine whether the classification was correct. The proportion correct for the test cases is then computed to quantify decoding accuracy.

Of course, we typically have far more than two electrode sites in a typical ERP study. All *N* sites are used simultaneously for decoding, giving us an *N*-dimensional space. If we have 30 electrode sites, for example, we will have a 30-dimensional space, and the SVM will determine the optimal *hyperplane* in this 30-dimensional space for distinguishing between the stimulus classes^2^. A separate SVM is trained and tested at each time point in the ERP waveform, yielding a waveform showing decoding accuracy at each time point (which is roughly analogous to an ERP difference wave because it shows the ability of the decoder to differentiate between the conditions).

Although the training and testing cases shown in Figure 1a are individual trials, single-trial EEG epochs are typically too noisy to allow good decoding of subtle stimulus classes. Instead, the trials for each class are typically divided into multiple sets of 10-20 trials, and an averaged ERP waveform is created from each of these sets. For example, if there were 80 face trials and 80 car trials, we might create five face averages and five car averages, each with 16 trials. This is illustrated in Figure 1b. We would train the SVM on the data from four face averages and four car averages and then test the SVM on the data from the one face average and the one car average that were not used for training (separately at each latency). This yields a smaller number of training and testing cases. However, SVMs are particularly well suited for learning to classify data on the basis of a small number of training cases, and the increase in signal-to-noise ratio produced by averaging outweighs the reduction in the number of training cases(Gholami & Fakhari, 2017; Grootswagers et al., 2017).

Many conventional ERP experiments also involve comparing two classes of trials^3^. For example, mismatch negativity (MMN) and P3b experiments typically involve comparing standard stimuli with oddball stimuli; N400 experiments typically involve comparing words that vary in terms of a factor such as semantic relatedness or concreteness; and error-related negativity experiments typically involve comparing trials with correct versus incorrect responses. In these experiments, an SVM can also be used to decode which of the two trial types occurred, and decoding accuracy is roughly analogous to the difference in voltage between the two trial types at a given latency.

Just as the difference in voltage is commonly averaged across a measurement window in conventional ERP analyses, decoding accuracy can be averaged across a measurement window. Decoding accuracy can therefore be used as the dependent variable in statistical analyses in exactly the same way as differences in voltage across experimental conditions (although the interpretation is subtly different, as will be described in the Discussion). However, because SVMs use the entire pattern of voltage over the scalp in an optimal manner for each individual participant, it is plausible that they might provide greater sensitivity, larger effect sizes, and improved statistical power relative to simple voltage differences.

Decoding accuracy is computed by taking a categorical variable (correct or incorrect) and aggregating across test cases to create a continuous variable (proportion correct). This is much like calculating proportion correct in a perception experiment. Unfortunately, such measures can have poor psychometric properties, such as an insensitivity to differences near the floor or ceiling. An alternative approach in perception experiments is to apply signal detection theory and compute the d’ metric of sensitivity, which represents the difference between the average percept produced by the two classes scaled by the within-class variability of the percept. In decoding, the analogous metric is the Mahalanobis distance, which is illustrated in an idealized manner in Figure 1c. This metric represents the distance between the centroids of the two single-trial distributions, scaled by the pooled multidimensional standard deviation and has previously been used in EEG research to decode differences in visual working memory states (Wolff et al., 2015). The Mahalanobis distance is also analogous to the Cohen’s *d* metric of effect size but extended to a multidimensional space. The Mahalanobis distance has no maximum value, so ceiling effects are not an issue.

However, whereas the chance value for decoding accuracy is usually very clear (e.g., 0.5 for a binary classification), chance is not well defined for the Mahalanobis distance (because distances can never be negative, and noise alone will cause the Mahalanobis distance to be greater than zero). This shortcoming can be solved with a cross-validation approach in which the data are split into random halves (see Method for more details). The result is the *cross-validated Mahalanobis distance* (or *crossnobis* distance; Walther et al., 2016).

The goal of the present study was to assess whether SVM-based decoding accuracy and the crossnobis distance might offer advantages for ERP studies in which the goal is to compare the amplitude of an ERP component across two conditions. Given that decoding can reliably determine which of two faces is being perceived or which of two orientations is being maintained in working memory, perhaps it could provide a more powerful way of asking whether, for example, the N170 component is larger for faces than for cars. To draw a conclusion that is relatively general, we analyzed data for seven different ERP components (N170, N2pc, P3b, N400, mismatch negativity, lateralized readiness potential, and error-related negativity) taken from the ERP CORE dataset (Kappenman et al., 2021). Each component in this dataset is isolated by examining the difference between two conditions (e.g., faces versus cars for the N170 component, error trials versus correct trials for the error-related negativity). In the standard univariate approach, the difference in amplitude between the two conditions is quantified at single electrode site or cluster of electrode sites. We compared this approach with SVM decoding accuracy and the crossnobis distance to see whether the multivariate approaches would be more sensitive to the difference between conditions. Other decoding approaches are also available, but a recent study found that SVM outperformed other methods in the context of the N400 component (Trammel et al., 2023).

The three different analysis methods we examined are on different scales (number of microvolts for the univariate analysis, proportion correct for SVM decoding, and multivariate distance in microvolts for crossnobis distance). To put them on a common scale so that we could make comparisons among them, we converted the set of values for the entire group of 40 participants to the Cohen’s *d_z_* metric of effect size (Cohen, 1988). This metric represents the size of an effect as the mean across participants minus chance scaled by the standard deviation across participants. Cohen’s *d_z_* is directly related to the statistical power for detecting a difference between experimental conditions in a within-subjects analysis, so a larger *d_z_* is preferable (all else being equal). Converting the results into *d_z_* values made it possible to compare the different analytic approaches on the same scale and allows for planning of future studies by other researchers (Lakens, 2013). Our primary question, therefore, was whether one or both multivariate techniques yields a larger *d_z_* than the traditional univariate approach.

All the decoding analyses reported here were carried out using tools that are built into versions 10 and higher of ERPLAB Toolbox (Lopez-Calderon & Luck, 2014). These tools allow the user to perform decoding using a graphical user interface without any programming (although we used scripts to automate the process and provide a record of exactly how the data were analyzed). Thus, it is now easy and inexpensive for anyone to apply decoding methods to their own ERP data.

## 2. Method

The ERP CORE (Kappenman et al., 2021) consists of stimulus presentation scripts, data, and data analysis scripts for six different experiments that can be used to isolate seven different ERP components. A complete description of the participants, stimuli, tasks, and preprocessing for the ERP CORE experiments can be found in the original paper. Here we provide a brief summary, along with a detailed description of the analyses that were performed for the present study.

### 2.2 Participants

Forty neurotypical young adults who provided informed consent performed each of the six ERP CORE experiments in a single session. For any given component, participants with EEG artifacts on a large number of trials (> 25% of epochs) were excluded. In addition, one participant had only two error trials for the ERN component and was excluded. These procedures led to the exclusion of 1 – 6 participants per component. Table 1 lists the final sample size for each component. The study was approved by the UC Davis Institutional Review Board.

**Table 1.**
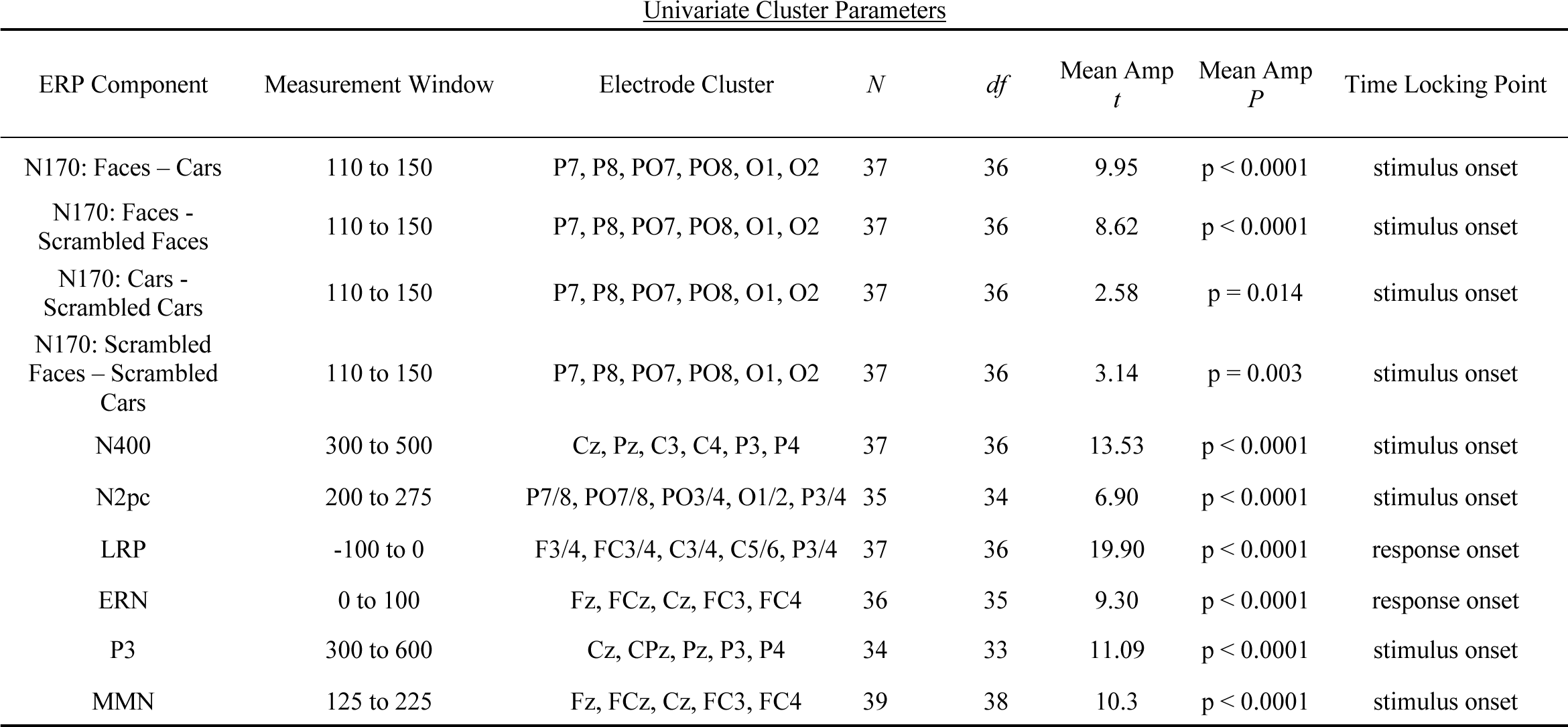
Univariate analysis parameters for each component, along with the corresponding *t* test.

| <u>Univariate Cluster Parameters</u> |  |  |  |  |  |  |  |
| --- | --- | --- | --- | --- | --- | --- | --- |
| ERP Component | Measurement Window | Electrode Cluster | $N$ | $df$ | Mean Amp<br>$t$ | Mean Amp<br>$P$ | Time Locking Point |
| N170: Faces – Cars | 110 to 150 | P7, P8, PO7, PO8, O1, O2 | 37 | 36 | 9.95 | $p < 0.0001$ | stimulus onset |
| N170: Faces - Scrambled Faces | 110 to 150 | P7, P8, PO7, PO8, O1, O2 | 37 | 36 | 8.62 | $p < 0.0001$ | stimulus onset |
| N170: Cars - Scrambled Cars | 110 to 150 | P7, P8, PO7, PO8, O1, O2 | 37 | 36 | 2.58 | $p = 0.014$ | stimulus onset |
| N170: Scrambled Faces – Scrambled Cars | 110 to 150 | P7, P8, PO7, PO8, O1, O2 | 37 | 36 | 3.14 | $p = 0.003$ | stimulus onset |
| N400 | 300 to 500 | Cz, Pz, C3, C4, P3, P4 | 37 | 36 | 13.53 | $p < 0.0001$ | stimulus onset |
| N2pc | 200 to 275 | P7/8, PO7/8, PO3/4, O1/2, P3/4 | 35 | 34 | 6.90 | $p < 0.0001$ | stimulus onset |
| LRP | -100 to 0 | F3/4, FC3/4, C3/4, C5/6, P3/4 | 37 | 36 | 19.90 | $p < 0.0001$ | response onset |
| ERN | 0 to 100 | Fz, FCz, Cz, FC3, FC4 | 36 | 35 | 9.30 | $p < 0.0001$ | response onset |
| P3 | 300 to 600 | Cz, CPz, Pz, P3, P4 | 34 | 33 | 11.09 | $p < 0.0001$ | stimulus onset |
| MMN | 125 to 225 | Fz, FCz, Cz, FC3, FC4 | 39 | 38 | 10.3 | $p < 0.0001$ | stimulus onset |

### 2.3 Overview of Stimuli and Tasks

Figure 2 illustrates the six experimental paradigms. In the N170 task (Rossion & Jacques, 2011), participants were shown faces, scrambled faces, cars, and scrambled cars (p = .25 for each category); they were instructed to indicate whether a given stimulus was intact or scrambled. In the N400 task, (Kutas & Federmeier, 2011), each trial consisted of a prime word followed by a semantically related or unrelated target word; participants were instructed to indicate whether each target word was related or unrelated to the preceding prime word. In the N2pc task, (Kiss et al., 2008), each stimulus array included a blue square, a pink square, and several black squares; participants were instructed to attend to the blue squares in some trial blocks and the pink squares in other trial blocks and indicate whether the gap on the attended-color square in each array was on the top or bottom of that square. In the flankers task used to elicit the lateralized readiness potential (LRP) and error-related negativity (ERN) (Eimer & Coles, 2003; Gehring et al., 2012) each display contained a left- or right-pointing central arrow flanked by distractor arrows that were pointing in the same direction or the opposite direction (with equal probability); participants were instructed to indicate the direction of the central arrow. To achieve an appropriate number of error trials for examining the ERN, feedback was used to encourage fast response times that produced incorrect responses on 10–20% of trials. In the P3b task, (Polich, 2012), the letters A, B, C, D, and E were presented in random order (p = .2 each); one letter was defined as the target at the beginning of each block, and participants were instructed to indicate whether each stimulus was the target letter (p = .2) or any of the nontarget letters (p = .8 for the nontarget category). In the MMN task (Näätänen et al., 2007), participants heard a randomized sequence of standard tones (1000 Hz, 80 dB, p = .8) and deviant tones (1000 Hz, 70 dB, p = .2); participants were instructed to ignore these tones and watch a silent video.

**Figure 2.**
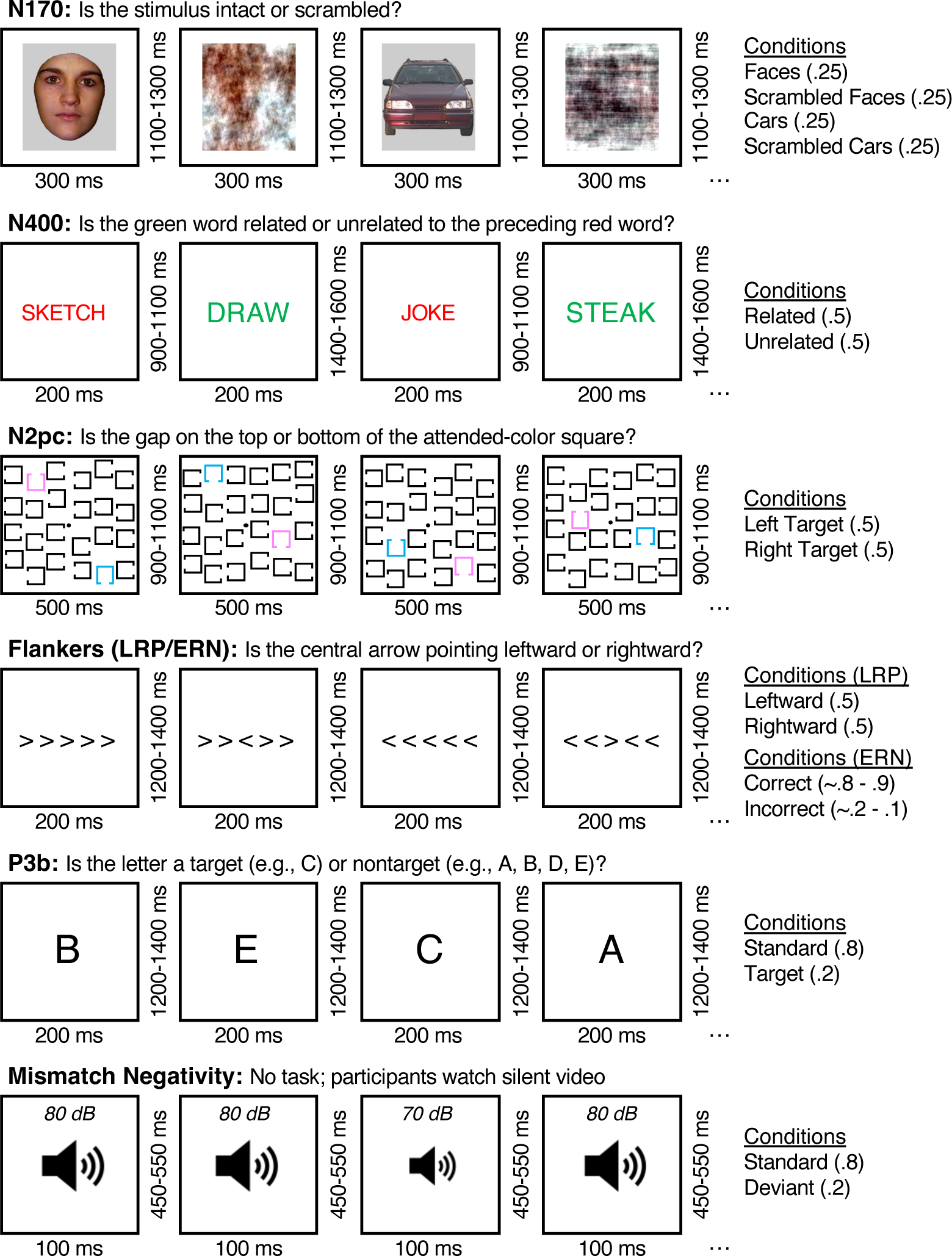
Overview of the ERP CORE tasks. In the N170 task, participants were presented with faces, scrambled faces, cars, and scrambled cars (p = .25 each) and were required to indicate whether each stimulus was intact or scrambled. In the N400 task, participants judged the semantic relatedness of each green target word to the preceding red prime word. The N2pc task involved attending to blue or pink squares in an array of black squares and indicating the location of the gap on the attended-color square. The flankers task was used to isolate both the lateralized readiness potential (LRP) and the error-related negativity (ERN). Participants were required to indicate the direction of a central arrow flanked by distractors. In the P3b task, participants were presented with a sequence of randomly ordered letters from the set {A, B, C, D, E} and were required to indicate whether each letter was a predefined target letter or one of the four nontarget letters. In the mismatch negativity (MMN) task, participants watched a silent video and were instructed to ignore a sequence of tones that included rare deviant tones (70 dB) and frequent standard tones (80 dB).

### 2.4 EEG Recording and Processing Procedures

All analyses described here began with the preprocessed, epoched data provided in the ERP CORE resource (Kappenman et al., 2021), which are freely available at https://doi.org/10.18115/D5JW4R. Continuous EEG recordings were obtained using a Biosemi ActiveTwo recording system with active electrodes placed at 30 scalp sites: FP1, F3, F7, FC3, C3, C5, P3, P7, P9, PO7, PO3, O1, Oz, Pz, CPz, FP2, Fz, F4, F8, FC4, FCz, Cz, C4, C6, P4, P8, P10, PO8, PO4, O2. Additional electrodes were placed lateral to the eyes and under the left eye to record the horizontal and vertical electrooculogram (HEOG and VEOG).

Ocular artifacts were corrected offline using independent component analysis, and trials were marked for exclusion if they contained behavioral errors, large voltage deflections in any channel, or ocular artifacts at the time of the stimulus. The data were high-pass filtered (non-causal Butterworth filter, half-amplitude cutoff at 0.1 Hz, 12 dB/octave), downsampled to 256 Hz, and segmented into 1000 ms epochs with 200 ms baselines. The data were referenced offline to the average of the P9 and P10 electrodes (near the left and right mastoids), except that the N170 data were referenced to the average of all scalp sites as is common in N170 studies^4^. See Kappenman et al. (2021) for extended details on the recording methods and preprocessing pipeline.

For the present analyses, we also applied a low-pass filter (non-causal Butterworth filter, half-amplitude cutoff at 20 Hz, 12 dB/octave) to the epoched EEG data prior to all further analyses which is commonly applied prior to measuring components. All preprocessing was conducted in MATLAB using the EEGLAB and ERPLAB toolboxes (Delorme & Makeig, 2004; Lopez-Calderon & Luck, 2014; MathWorks, 2023).

### 2.5 Traditional Univariate Analyses

The traditional univariate analysis began by computing a difference wave between the key conditions for a given ERP component for each participant (e.g., oddballs versus standards for the P3b component). We then averaged the difference waves across an optimal cluster of electrode sites for that component, as determined by Zhang & Kappenman (2023). An alternative approach would have been to use the maximal electrode site where a component is known to be represented (e.g., site Pz for the P3b) but when we did these univariate analyses, we found that using a cluster of electrodes led to very similar results (see supplementary Figures S2-S4). We therefore chose to use clusters of electrode sites as Zhang & Kappenman (2023) showed cluster-based measurements are as good and often better than measuring from single sites. We then measured the mean voltage across an a priori measurement window and used a one-sample *t* test to determine whether this difference was significantly different from zero (which is equivalent to asking whether the two conditions were significantly different from each other). Given that the direction of the effects in these paradigms is already known from hundreds of previous studies for a given component, these tests were one-tailed. The alpha for these and all other statistical tests was 0.05. The measurement windows and electrode sites for each component are shown in Table 1. Supplementary Table S1 shows the analogous results when the data were obtained from the single electrode where the component is largest.

The effect size for the mean amplitude across the measurement window or for an individual time point was quantified with Cohen’s *d_z_*. This was computed from the voltage in the difference waves (e.g., faces minus cars for N170, contralateral minus ipsilateral for N2pc and LRP), which is identical to computing the effect size comparing the parent waves. For difference scores such as these, the Cohen’s *d_z_* equation is:

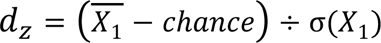

where X_1_ is equal to the mean difference wave voltage across participants, chance is set to 0 meaning no difference between conditions, and σ(X_1_) is the standard deviation of the difference wave voltages across participants (see Lakens, 2013).

We also conducted a mass univariate analysis (Groppe et al., 2011; Maris & Oostenveld, 2007) in which we performed a one-sample *t* test comparing the difference score to zero at each time point between 0 ms and the end of the epoch, using the false discovery rate (FDR) correction for multiple comparisons (Benjamini & Yekutieli, 2001). Given that the direction of the effects in these paradigms is already known from hundreds of previous studies for a given component, one-tailed tests were performed (as shown with gray shading in the waveform figures). However, the direction of these one-tailed tests was based on a specific component that occurs during a particular time window (e.g., the N170 time window in our comparison of faces versus cars). Effects of the opposite polarity were sometimes obtained outside this time window (e.g., a positivity that followed the N170), which were not significant by definition with the one-tailed test. We therefore also conducted two-tailed statistical tests (show with horizontal black lines in the waveform figures).

Because Cohen’s *d_z_* is negative when the voltage is more negative in one condition than in another, we took the absolute value of the effect size. The standard error and 95% confidence interval of each d_z_ value was estimated using bootstrapping, as described in Section 2.8.

### 2.6 Support Vector Machine (SVM) Decoding Analyses

We performed SVM decoding using the decoding tools built into ERPLAB Toolbox, which rely on Matlab’s fitcsvm() function. For each component, the two conditions for that component served as the classes being decoded (e.g., faces vs. cars for the prototypical N170 comparison). Decoding was conducted separately at each time point for each participant. Thus, we obtained a waveform for each participant that reflected the decoder’s ability to distinguish between the two conditions at each time point, which is the MVPA analog of the difference in voltage between the two conditions at each time point (i.e., the difference wave).

The SVM analyses used an *N*-fold cross-validation procedure. Specifically, we randomly divided the data for a given condition into *N* sets of trials and created an average ERP from each of those N sets of trials. For a given component, we selected the largest *N* that would yield 10–20 trials per average. For example, there were 80 face trials and 80 car trials for the N170, and we used five sets of averages (five *crossfolds*) for each condition, with 16 trials per average prior to the exclusion of trials with artifacts or errors. Table 2 shows the number of crossfolds and the minimum number of trials remaining in each condition after exclusion for each component.

**Table 2.**
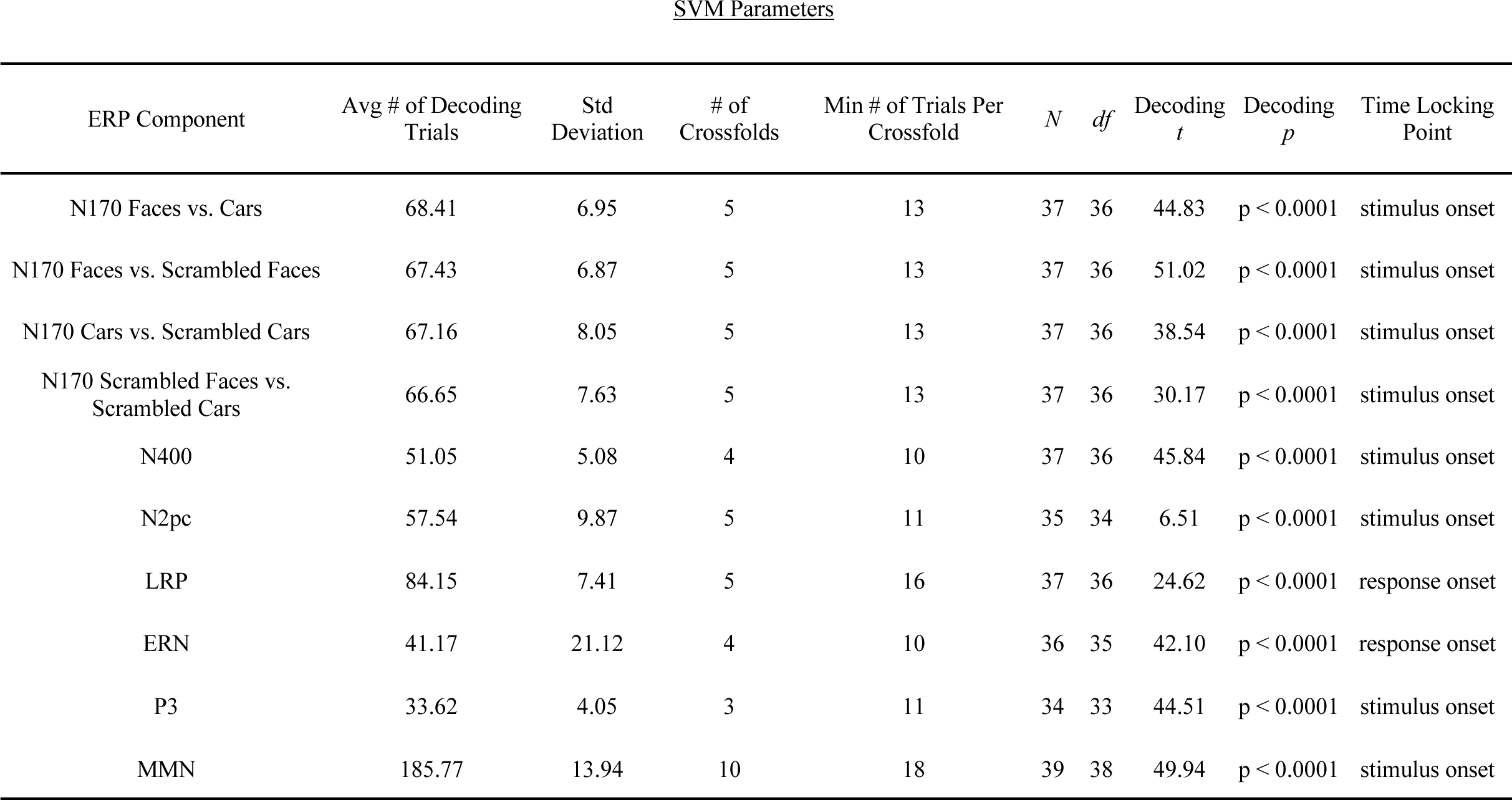
Decoding parameters for all analyses done on each component. The number of crossfolds was determined so that 10-18 trials were included in the averaged ERPs for a given component. The *t* values shown here represent one-sample *t* tests against chance for the average decoding accuracy across the measurement window for a given component.

| ERP Component | Avg # of Decoding Trials | Std Deviation | # of Crossfolds | Min # of Trials Per Crossfold | $N$ | $df$ | Decoding $t$ | Decoding $p$ | Time Locking Point |
| --- | --- | --- | --- | --- | --- | --- | --- | --- | --- |
| N170 Faces vs. Cars | 68.41 | 6.95 | 5 | 13 | 37 | 36 | 44.83 | $p < 0.0001$ | stimulus onset |
| N170 Faces vs. Scrambled Faces | 67.43 | 6.87 | 5 | 13 | 37 | 36 | 51.02 | $p < 0.0001$ | stimulus onset |
| N170 Cars vs. Scrambled Cars | 67.16 | 8.05 | 5 | 13 | 37 | 36 | 38.54 | $p < 0.0001$ | stimulus onset |
| N170 Scrambled Faces vs. Scrambled Cars | 66.65 | 7.63 | 5 | 13 | 37 | 36 | 30.17 | $p < 0.0001$ | stimulus onset |
| N400 | 51.05 | 5.08 | 4 | 10 | 37 | 36 | 45.84 | $p < 0.0001$ | stimulus onset |
| N2pc | 57.54 | 9.87 | 5 | 11 | 35 | 34 | 6.51 | $p < 0.0001$ | stimulus onset |
| LRP | 84.15 | 7.41 | 5 | 16 | 37 | 36 | 24.62 | $p < 0.0001$ | response onset |
| ERN | 41.17 | 21.12 | 4 | 10 | 36 | 35 | 42.10 | $p < 0.0001$ | response onset |
| P3 | 33.62 | 4.05 | 3 | 11 | 34 | 33 | 44.51 | $p < 0.0001$ | stimulus onset |
| MMN | 185.77 | 13.94 | 10 | 18 | 39 | 38 | 49.94 | $p < 0.0001$ | stimulus onset |

Because spurious above-chance decoding accuracy is likely if the number of trials differs between the conditions being decoded for a given participant, we equated (*floored*) the number of trials used for decoding in each participant. That is, we determined which condition had the smallest number of trials for a given participant after exclusion of trials with errors and artifacts, and we randomly selected an equivalent number of trials for the other condition for that participant. This was done separately for each component. For components with very different numbers of trials per condition (P3b, MMN, and ERN), this meant that a relatively small subset of the trials was selected from the condition with the larger number of trials. For example, there were 160 frequent trials and 40 rare trials for the P3b component, and we used only 40 of the frequent trials (or even fewer given the exclusion of trials with errors and artifacts)^5^.

We performed decoding on the *N* averaged ERP waveforms for each condition, separately at each time point. We included only the 28 scalp EEG channels, excluding artifact channels and reference channels. The decoding was performed once for each of the *N* crossfolds, using *N*-1 of the averaged ERPs from each condition for training and the remaining 1 averaged ERP for each condition for testing. For each of these crossfolds, the SVM was trained using the pattern of voltages across the 28 electrode sites at the time point of interest from each of the *N*-1 averages for each of the two conditions (e.g., from 4 averages of face trials and 4 averages of car trials). From these training examples, the SVM determined an optimal classification hyperplane for that time point. The SVM was then tested by being given the pattern of voltage across electrodes at that time point for each of the two averages that were not used for training (e.g., 1 average of face trials and 1 average of car trials) and asked to guess the condition for each of these patterns (e.g., face or car). Thus, for a given crossfold, the SVM made two binary guesses, one for each condition. Because there were 2 test cases and therefore 2 guesses for each crossfold, there were 2*N* test cases and 2*N* guesses across the *N* crossfolds (e.g., 10 total guesses across 5 crossfolds for the N170 *faces/cars* component). This process was then repeated for each time point.

To increase the resolution of the decoding accuracy, we repeated this process 100 times for each participant (100 *iterations*). For each iteration, we re-randomized the assignment of trials to averages and trained new SVMs. This gave us 200*N* guesses for a given component at each time point for each fold (e.g., 1000 total guesses for the 5-fold N170 analysis). Decoding accuracy for a given time point was computed as the proportion of guesses that were correct across these guesses. Because there were two conditions for a given component, and the number of training cases was always the same for the two conditions, chance decoding accuracy was 0.5 at each time point.

For a given component, this process ultimately yielded a decoding accuracy value at each time point for each participant. To determine whether decoding accuracy at each time point was significantly above chance for the group of participants, we conducted a mass univariate analysis using a separate one-sample *t* test against chance (0.5) at each time point, with the FDR correction for multiple comparisons (Benjamini & Yekutieli, 2001). Although we computed decoding accuracy for each point in the entire epoch, we excluded the prestimulus period from the FDR correction. One-tailed tests were used because below-chance accuracy is not meaningful with this procedure^6^. We also computed the Cohen’s *d_z_* metric of effect size:

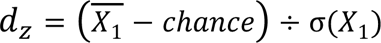

where X_1_ is equal to the mean of the decoding accuracy values across participants, chance is 0.5 or 50%, and σ(X_1_) is the standard deviation of the decoding accuracy values across participants. This is exactly the same d_z_ formula used for the univariate analyses. As detailed in Section 2.8, bootstrapping was used to compute standard errors and 95% confidence intervals for *d_z_*.

In many ERP studies, the amplitude of a component is quantified as the mean voltage within a specific time window. To conduct an analogous analysis with the decoding data, we simply averaged the decoding accuracy values across the time points within the measurement window listed in Table 2 for a given component (e.g., 110-150 ms for Face vs Cars). A different approach would be to average the voltage across the measurement window prior to decoding, but we found that averaging across time points after decoding tended to yield larger effect sizes (see supplementary Figure S1).

The univariate analyses used data only from the cluster of electrode sites where the component was known to be largest, whereas the SVM analyses used all of the electrode sites, even those at which the component was near zero. We tested the possibility that this might give the univariate analyses an advantage. Specifically, we repeated the SVM analyses using only the electrode sites used for the univariate analyses (listed in Table 1). We found that this approach led to similar or worse results than using all of the electrode sites for decoding (see supplementary Figures S5–S7 and Table S2). We therefore used all of the electrode sites for the decoding analyses reported here.

### 2.7 Crossnobis Analyses

As illustrated in Figure 1C, the Mahalanobis distance is defined as the distance between the centroids of the two classes, scaled by the multidimensional standard deviation of each class. Whereas SVMs rely on good estimates of the values near the decision line to work well, which is facilitated by averaging, individual data points have less impact on the Mahalanobis distance. However, the Mahalanobis distance benefits from using a large number of data points to obtain robust estimates of the multidimensional standard deviation of a given class, so single-trial data points are typically used instead of averaged ERPs. Because averaging is unnecessary, there is also no need to equate the number of trials in the two conditions being compared, so all the data can be used.

Figure 1c shows a simplified example of crossnobis distance in which ERP data from two electrode sites (e.g., PO8 and Cz) at a particular time point (e.g., 170 ms after stimulus onset) are used to decode whether the subject had been shown a picture of a face or car. The Mahalanobis distance at a given time point is calculated as:

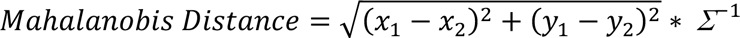

where x and y correspond to the mean voltages at the two electrodes, the subscripts 1 and 2 correspond to the two conditions (e.g., faces and cars), and Λ corresponds to the multi-dimensional standard deviation across all observations. Note that the centroids and multidimensional standard deviations are computed using exactly the same approach used in linear discriminant analysis.

One shortcoming of the Mahalanobis distance is that it is always positive, so chance is not zero and is difficult to determine. This can be seen in the equation, where the differences between condition means (*x_1_* - *x_2_* and *y_1_* - *y_2_*) are both squared, creating positive values. This shortcoming can be solved by using the crossvalidated Mahalanobis distance (or *crossnobis distance*; Walther et al., 2016), which involves dividing the data into two evenly random splits, *A* and *B*, for each condition. This updated metric for a given time point is then:

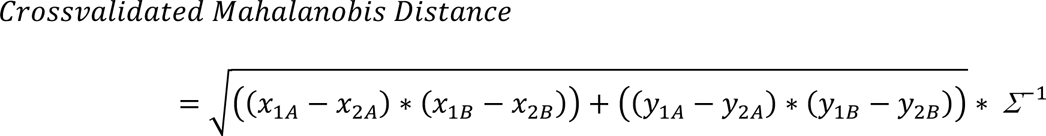

where the equation is the same as used for the Mahalanobis distance, except rather than computing a single squared difference score across conditions at an electrode site (e.g., (*x*_1_ – *x*_2_)^2^), two independent difference scores are computed across the subsets (*A* and *B*) of data across conditions at an electrode site and then multiplied (i.e., ((*x*_1*A*_ – *x*_2*A*_) ∗ (*x*_1*B*_ – *x*_2*B*_))). Because there is no squaring of values^7^, the cross-validated distance can be either positive or negative, and the expected chance value is zero (i.e., if the centroids of the two conditions do not differ). Thus, if the crossnobis distance is significantly different from zero, one can conclude that the two conditions are different.

The cross-validation procedure was repeated 100 times with different random splits of the data to provide a more robust result. Each iteration produced a distance value, and we simply averaged these distance values across iterations. Note that, unlike SVM decoding accuracy, the crossnobis distance is not affected by differences in the number of trials in each class. Consequently, we used all trials except for those with behavioral errors or artifacts.

Like the decoding analysis, this analysis was performed separately at each time point for a given component, yielding a waveform of distance values across time points for each participant. As for the SVM decoding, we assessed statistical significance via a one-sample *t* test against chance (zero) at each time point (excluding the prestimulus period), with an FDR correction for multiple comparisons (Benjamini & Yekutieli, 2001). We also averaged the single-point crossnobis distances over the recommended measurement window for a given component and compared the result to chance (zero) with a one-sample *t* test. Negative crossnobis values are not meaningful with the present procedure, so these tests were one-tailed. We also attempted a variant of this analysis in which we averaged the timepoints into a mean amplitude prior to computing the crossnobis distance. However, this approach was consistently worse than our original approach (see supplementary Figure S1). Additionally, we attempted this analysis using only the set of electrodes used for the univariate analysis of a given component, but this did not produce a clear advantage over using all electrode sites (see supplementary Figures S8–S10 and Table S3), so we chose to use all sites in the primary analyses. Finally, we computed the effect size (*d_z_*) of the crossnobis distance:

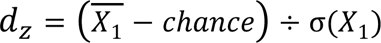

where X_1_ is equal to the mean of the distance values across participants, chance is 0, and σ(X_1_) is the standard deviation of the distance values across participants. This is exactly the same d_z_ formula used for the univariate and SVM analyses. Bootstrapping was used to compute standard errors and 95% confidence intervals for *d_z_*, as detailed in the next section.

### 2.8 Bootstrapping Procedures

We used a standard bootstrapping approach (Efron & Tibshirani, 1994) to estimate the standard error of the effect size values, both at individual time points and when averaged across the measurement window for a given component. For a given score (i.e., a voltage in a univariate difference wave, a cronssnobis value, or a decoding accuracy value), we selected a random sample of *N* of the *N* participants (sampling from the set of *N* participants with replacement) and computed the effect size for this random sample. We iterated this process 10,000 times with different random samples, obtaining an effect size for each iteration, and the standard error was defined as the standard deviation of these 10,000 effect sizes. For the point-by-point analyses, this was done separately at each time point to obtain a standard error at each time point. For the time-window analyses, the scores were averaged across the time window prior to computing the effect size.

We also used bootstrapping to compare the effect sizes between the three analysis methods (univariate, crossnobis, and SVM). Specifically, we computed the percentage difference in effect size between each pair of analysis methods. The percentage difference was defined as 100 × (*A* – *B*) / *B*, where *A* and *B* represent the effect sizes for two different methods (e.g., SVM decoding accuracy versus univariate difference voltage). We used bootstrapping to estimate the 95% confidence interval for a given percentage difference. For each of 10,000 bootstrapping iterations, we obtained the percentage difference from a random sample of *N* of the *N* participants (sampled with replacement), yielding 10,000 estimates of the percentage difference. In idealized data, the bounds of the 95% confidence interval would be defined as the 2.5 and 97.5 percentiles of this set of estimates, but this can lead to bias in real data, especially if the distribution of values is asymmetrical. We therefore used the 95^th^ percentile *bias-corrected and accelerated* confidence intervals (Diciccio & Efron, 1996), as implemented in R using the package boot (Canty & Ripley, 2022; Davison & Hinkley, 1997b; R Core Team, 2023).

## 3. Results

We begin the description of the results with analyses of the N170 experiment, which included four different comparisons: faces versus cars, faces versus scrambled faces, cars versus scrambled cars, and scrambled faces versus scrambled cars. This is followed by an analysis of the N400, N2pc, LRP, and ERN components, each of which involved only two conditions. Those analyses are followed by analyses of the MMN and P3b components, which were obtained in oddball paradigms and therefore involved comparing conditions with very different numbers of trials.

### 3.1 N170 Point-by-Point Results

Figure 3 shows the N170 results when analyzed at each individual time point (the *point-by-point* analysis). The left column shows the traditional univariate analysis (the grand average ERP difference wave at the average of the electrode sites listed in Table 1). The classic comparison of faces and cars (Figure 3A) yielded the expected finding of greater N170 amplitude for faces than for cars. This effect was statistically significant from 100-150 ms (after correction for multiple comparisons). Small but significant differences were also present at other times (indicated by the gray shading in Figure 3). Intact faces and cars yielded a significantly more negative voltage than scrambled faces and cars from approximately 100-600 ms (Figures 3B and 3C). Interestingly, scrambled faces produced a significantly more positive voltage than scrambled cars from 136-200 and 230-292 ms; to our knowledge, this effect has not previously been reported.

**Figure 3.**
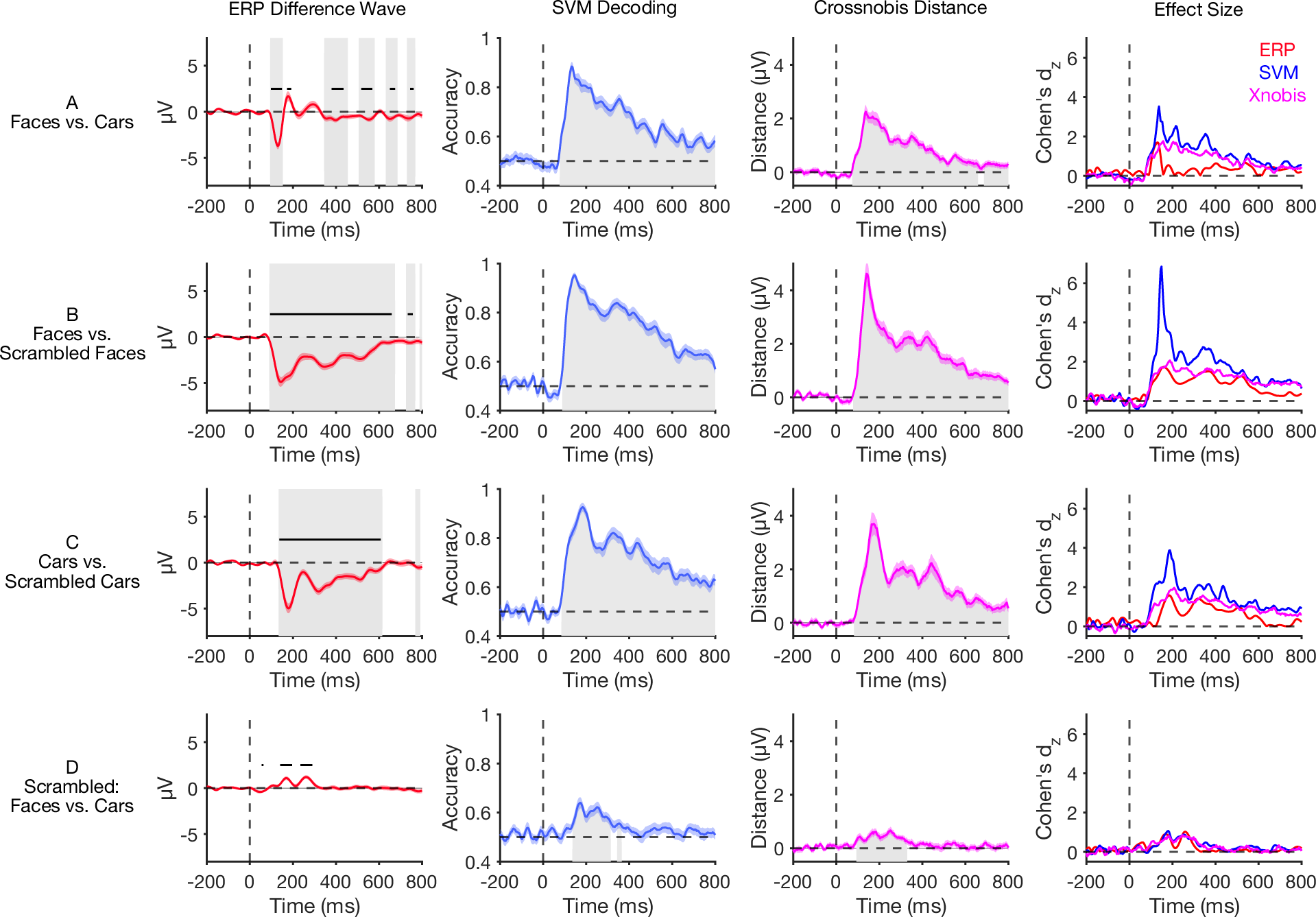
Point-by-point results for the N170 task. The four columns show the mean ERP difference voltages (averaged across the cluster of electrodes listed in Table 1), mean SVM decoding accuracy values, mean crossnobis distances, and effect sizes at each time point. The shaded bands around a given waveform show the standard error of the mean. Gray shading indicates time points at which the value is significantly different from chance with a one-tailed test in the direction of the expected effect after correction for multiple comparisons. The horizontal lines for the ERP difference waves indicate time periods at which the value is significantly different from chance with a two-tailed test (which can reveal effects with an unexpected polarity).

The second column of Figure 3 shows point-by-point SVM decoding accuracy, averaged across participants. Decoding accuracy was significantly greater than chance across a broader range of time points compared to the traditional univariate analysis of the difference waves. For example, face versus car decoding was significantly above chance from approximately 70 ms post-stimulus until the end of the epoch, whereas the univariate difference between faces and cars did not reach significance until 100 ms post-stimulus and was significant only during a small set of brief time periods. It should be noted, however, that the univariate ERPs at other electrode sites did differentiate between faces and cars at times when the ERPs from the electrodes shown in Figure 3A did not, so the long-lasting decoding is not incompatible with the ERP data. Similarly, the univariate difference between scrambled faces and scrambled cars (Figure 3D) was statistically significant in two brief periods, whereas a sustained significant decoding effect was present from 136-312 ms and from 343-363 ms.

The third column of Figure 3 shows the crossnobis distance results, which contained broad periods of statistical significance (93-339 ms) like those obtained for SVM decoding. Note that crossnobis distance is in units of microvolts because it is a distance between the voltage distributions for the conditions being contrasted. However, this metric is scaled by the multivariate standard deviation, so the crossnobis distances are not directly comparable to the univariate ERP amplitudes (even though both are in units of microvolts). Decoding accuracy is expressed in proportion correct, which is even more difficult to relate to the units of the univariate voltage difference and the crossnobis distance. Statistical significance is in the same units for all three analysis methods, but significance is a categorical variable and therefore not a very sensitive metric for comparing the three methods. In addition, one-tailed *t* test were performed for the decoding measures whereas two-tailed *t* tests were needed for the univariate voltage measure.

To directly compare the three analysis methods, we converted each of them into the effect size (Cohen’s d_z_) at each time point. This metric is continuous rather than categorical, and it is directly related to statistical power (Cohen, 1988). The effect sizes are shown in the rightmost column of Figure 3. We took the absolute value of each *d_z_* value for the univariate voltage so that those effect sizes would be positive whether the difference wave contained a positive voltage (e.g., the P3b effect) or a negative voltage (e.g., the N400 effect). The multivariate methods produced positive values for a true effect regardless of the polarity of the effect, so we did not take the absolute value for these methods.

In most cases, the effect size was largest for SVM decoding accuracy, which was more than three times as large as the effect size for the univariate ERPs in some cases (e.g., from 133-172 ms for the faces vs. scrambled faces comparison). The crossnobis effect size tended to be smaller than the SVM decoding effect size but larger than the univariate effect size. The one exception was the comparison of scrambled faces and scrambled cars, for which the effect sizes were similar across the three analysis methods. The general conclusion from these point-by-point analyses is that the multivariate techniques were at least as sensitive as the traditional univariate approach and often considerably more sensitive.

Although univariate analyses sometimes examine point-by-point differences between conditions, as in mass univariate analyses (Groppe et al., 2011; Maris & Oostenveld, 2007), it is more common for univariate analyses to examine the mean voltage across a measurement window. The next set of analyses compared this *time-window univariate approach* to decoding and crossnobis values averaged across the same time window.

### 3.2 N170 Time-Window Results

Figure 4 shows the results from the N170 paradigm, averaged across a time window of 110–150 ms (which was the time window recommended for the N170 component; Kappenman et al., 2021). Each of these effects was significantly different from chance (see Tables 1-3 for individual statistical tests). The effect size for SVM decoding accuracy was 2–3 times as large as the effect size for the univariate voltage difference for the comparisons of faces versus cars (Figure 4A), faces versus scrambled faces (Figure 4B), and cars versus scrambled cars (Figure 4C). The effect size for crossnobis distance was substantially smaller than the effect size for SVM decoding accuracy in these three cases but was as large or larger than the univariate effect size. As was observed in the point-by-point analyses (Figure 3), the effect sizes were similar across the three analysis methods for the comparison of scrambled faces and scrambled cars (Figure 4D), although the effect size was slightly larger for the crossnobis multivariate method than for the univariate method.

**Figure 4.**
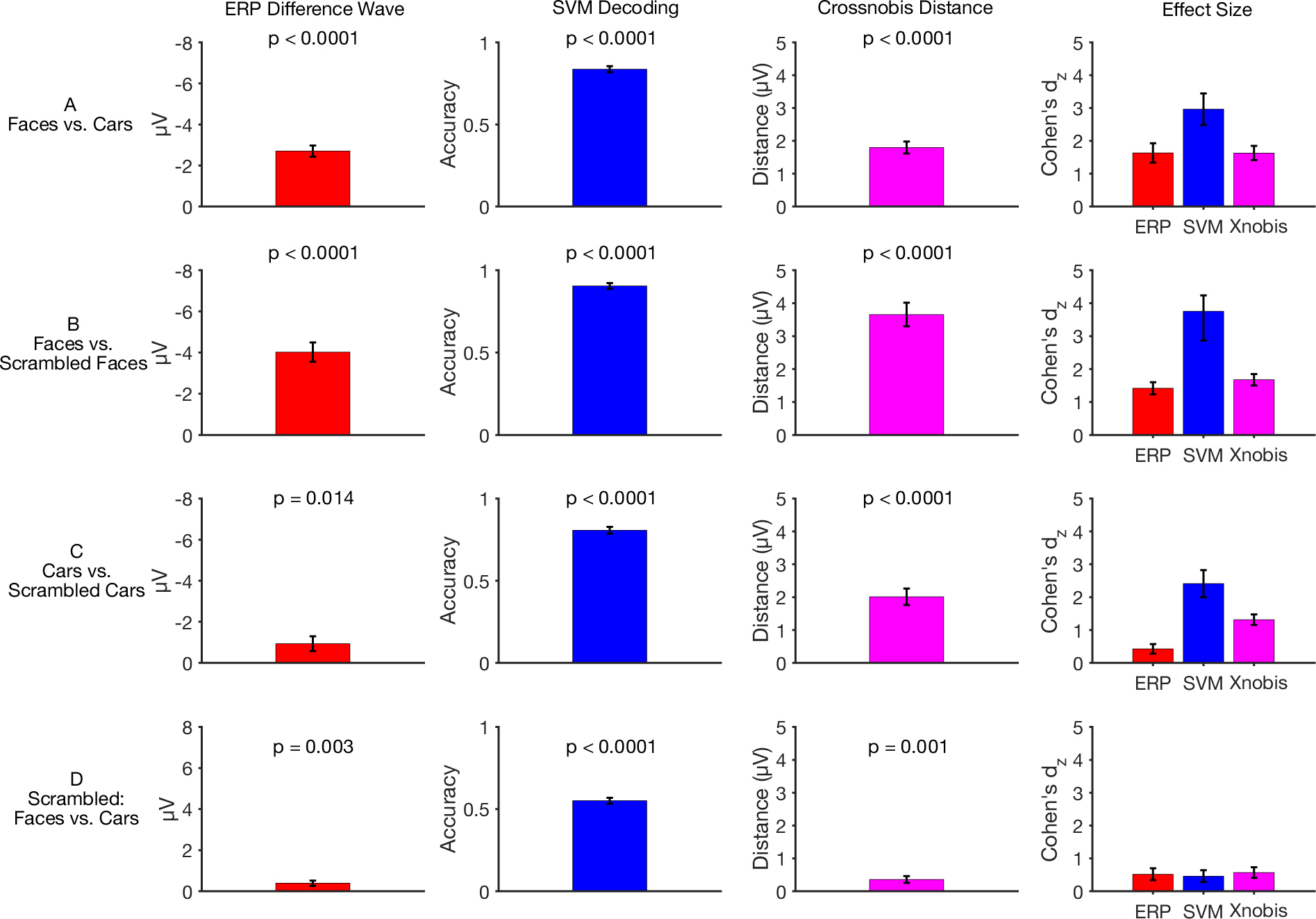
Time-window univariate ERP difference voltage, SVM decoding accuracy, and crossnobis distance, averaged across participants, for the four key comparisons in the N170 task. The time window was 110-150 ms for all three analysis methods. The p values are from one-tailed tests against chance. The right column shows the effect size (Cohen’s *d_z_*) for each analysis method. Error bars show ±1 SEM.

**Table 3.** Crossnobis parameters used for each individual component. Here the t-stats are more comparable to the univariate results. There is no indication of number of trials because this technique, like the univariate approach, allows researchers to use all trials.

| ERP Component | *N* | *df* | Xnobis *t* | Xnobis *p* | Time Locking Point |
| --- | --- | --- | --- | --- | --- |
| N170 Faces vs. Cars | 37 | 36 | 9.91 | $p < 0.0001$ | stimulus onset |
| N170 Faces vs. Scrambled Faces | 37 | 36 | 10.19 | $p < 0.0001$ | stimulus onset |
| N170 Cars vs. Scrambled Cars | 37 | 36 | 7.99 | $p < 0.0001$ | stimulus onset |
| N170 Scrambled: Faces vs. Cars | 37 | 36 | 3.46 | $p = 0.001$ | stimulus onset |
| N400 | 37 | 36 | 9.82 | $p < 0.0001$ | stimulus onset |
| N2pc | 35 | 34 | 4.14 | $p = 0.0002$ | stimulus onset |
| LRP | 37 | 36 | 14.15 | $p < 0.0001$ | response onset |
| ERN | 36 | 35 | 9.76 | $p < 0.0001$ | response onset |
| P3 | 34 | 33 | 8.59 | $p < 0.0001$ | stimulus onset |
| MMN | 39 | 38 | 7.08 | $p < 0.0001$ | stimulus onset |

### 3.3 N400, N2pc, LRP, and ERN Results

Figure 5 shows the point-by-point results for the N400, N2pc, LRP, and ERN components, and Figure 6 shows the results averaged over the recommended measurement window for each component.

**Figure 5.**
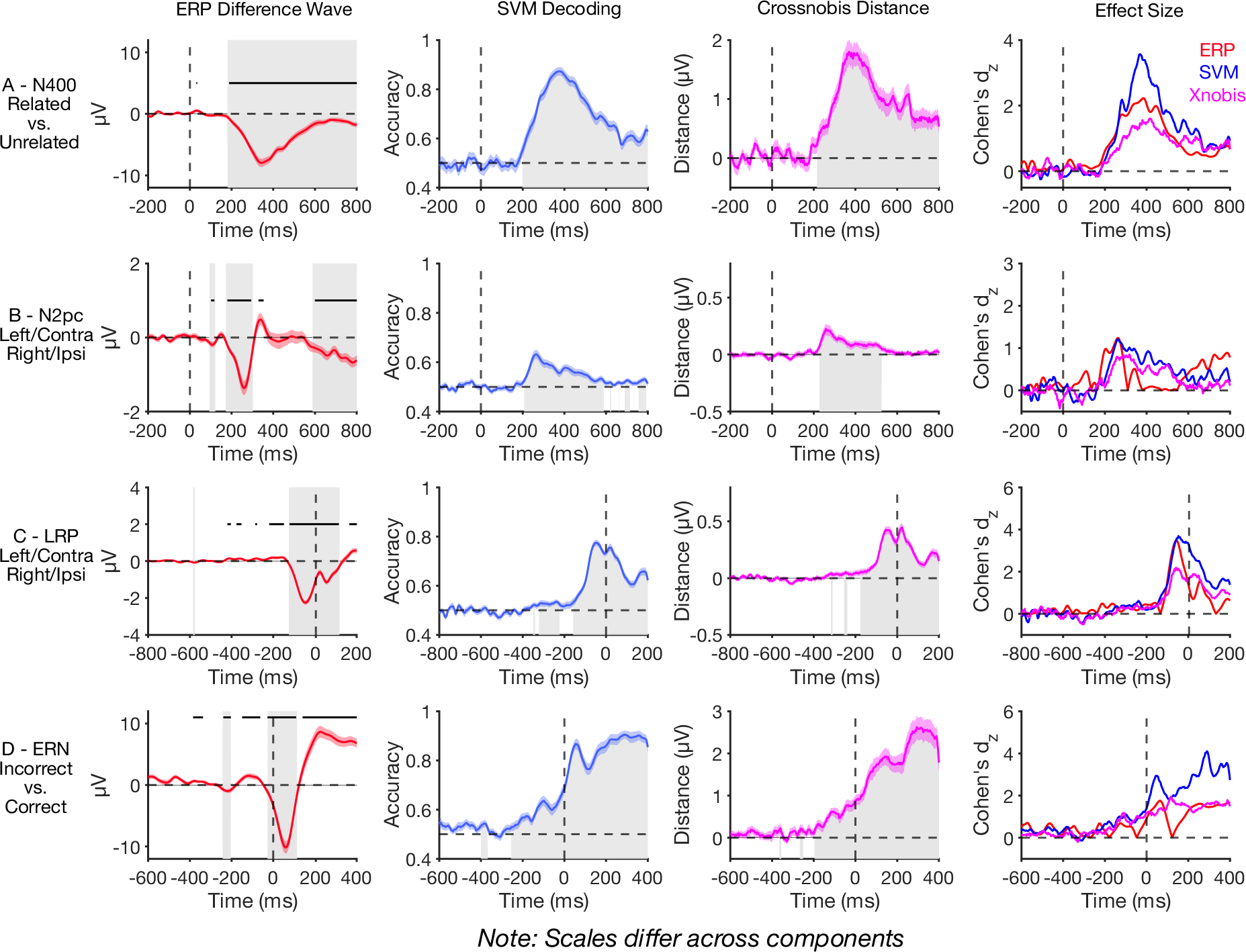
Point-by-point results for the N400, N2pc, LRP, and ERN components. The four columns show the mean ERP difference voltages (averaged across the cluster of electrodes listed in Table 1), mean decoding accuracy values, mean crossnobis distances, and effect sizes at each time point. The shaded bands around the a given waveform show the standard error of the mean. Gray shading indicates time points at which the value is significantly different from chance with a one-tailed test in the direction of the expected effect after correction for multiple comparisons. The horizontal lines for the ERP difference waves indicate time periods at which the value is significantly different from chance with a two-tailed test (which can reveal effects with an unexpected polarity).

**Figure 6.**
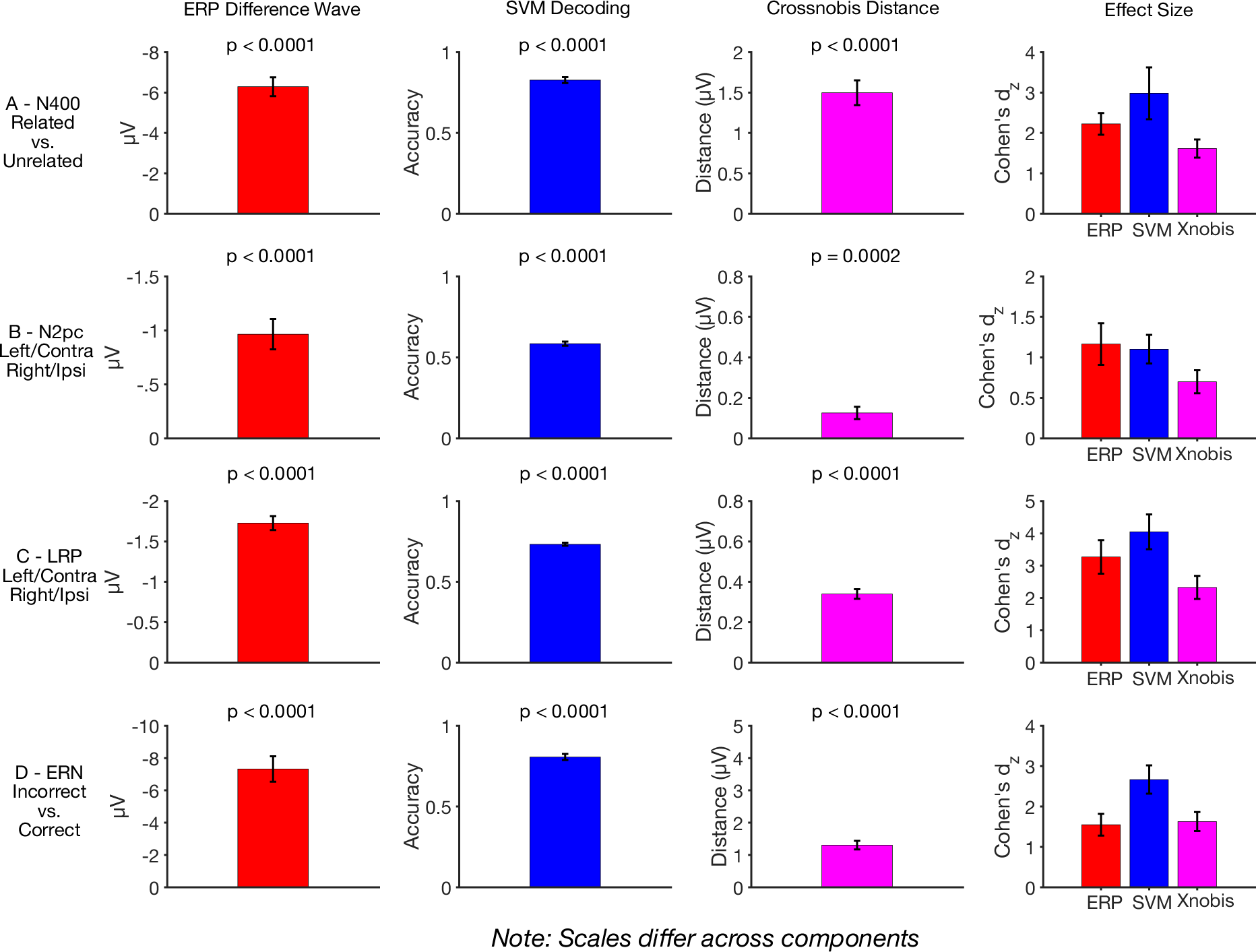
Time-window univariate ERP difference voltage, SVM decoding accuracy, and crossnobis distance, averaged across participants, for the N400, N2pc, LRP, and ERN components. The measurement window was 300 to 500 ms relative to the stimulus for the N400, 200 to 275 ms relative to the stimulus for the N2pc, -125 to 0 ms relative to the response for the LRP, and 0 to 100 ms relative to the response for the ERN. The *p* values are from one-tailed tests against chance. The right column shows the effect size (Cohen’s *d_z_*) for each analysis method. Error bars show ±1 SEM.

### 3.4 N400

In the N400 data (Figures 5A and 6A), the univariate comparison of related versus unrelated words yielded a prototypical negative N400 deflection that became statistically significant at 200 ms, peaked at approximately 400 ms, and remained significant throughout the remainder of the epoch. The SVM decoding and crossnobis distance analyses also sought to distinguish between related and unrelated words and yielded a similar time course. However, the effect sizes shown in the rightmost column of Figure 5A indicate that the effect size was much larger for SVM decoding accuracy than for the univariate voltage difference. By contrast, the effect size was smaller for the crossnobis distance than for the univariate voltage difference. The same pattern of effect sizes was obtained when the data were averaged across the N400 measurement window (300 to 500 ms; see Figure 6A).

### 3.5 N2pc

In the N2pc paradigm (Figure 5B), the contralateral-minus-ipsilateral difference wave yielded the typical pattern of a negative N2pc that was significant from 175-300 ms, followed by a brief post-N2pc positivity that was significant from 324-355 ms and then a sustained posterior contralateral negativity (SPCN; Kappenman & Luck, 2012; Mazza et al., 2007; Prime & Jolicoeur, 2010) that was significant from 593-800 ms. For the two multivariate methods, we decoded whether the target was contralateral or ipsilateral to the electrode sites. The midline sites were not used (Fz, FCz, Cz, CPz, Pz, Oz).

For SVM decoding, we first created averages of pairs of trials, one trial with the target in the left visual field (LVF) and one trial with the target in the right visual field (RVF). We averaged the left hemisphere channels for the RVF target trial with the right hemisphere channels for the LVF target trial to create a set of contralateral channels, and we averaged the left hemisphere channels for the LVF target trial with the right hemisphere channels for the RVF target trial to create a set of ipsilateral channels. We then performed the SVM decoding procedure described in Section 2.6, using contralateral and ipsilateral as the classes being decoded.

The crossnobis analysis also used these contralateral and ipsilateral classes, but we needed to ensure that different trials were used for the two splits of the data to avoid artificially minimizing the difference between the two splits. Specifically, the splits were defined by the odd and even trials. In all other respects, the analysis was the same as described in section 2.7. We did this for each color independently to avoid any possible sensory biases effecting the decoding, and then collapsed across the colors.

SVM decoding accuracy became statistically significant at 210 ms, which was slightly later than the univariate effect, and then remained significant until 585 ms. There were also brief periods of significance toward the end of the epoch, from 687-710 ms and from 753-789 ms. The crossnobis distance became significant at 226 ms and remained significant until 523 ms. The effect size (rightmost column of Figure 5B) was larger for the univariate analysis than for the multivariate analyses from 160-265 ms, after which it was larger for the SVM analyses from approximately 265-585 ms, and then became larger for the univariate analysis again during the SPCN period (from approximately 600 ms through the end of the epoch). The crossnobis analyses was larger than the N2pc analyses from 280-562 ms but not larger than the SVM. When the data were averaged across the N2pc measurement window (200 to 275 ms), the effect size was slightly larger for the univariate voltage than for the two multivariate methods (Figure 6B).

Note that the univariate ERP waveform contained a sequence of a negative N2pc followed by a post-N2pc positivity, which are thought to reflect two distinct neurocognitive processes (Gaspelin et al., 2023). However, there was a single period spanning these two components in which the SVM decoding accuracy and crossnobis distance was significantly greater from chance. This illustrates a potential downside of using multivariate pattern analysis methods to analyze ERP data, namely the loss of information about different polarities that might indicate different ERP components.

### 3.6 LRP

In the flankers paradigm, the response-locked LRP analysis (Figure 5C) yielded a negative voltage in the contralateral-minus-ipsilateral difference wave that began approximately 128 ms prior to the response and lasted until 113 ms after the response, at which point the voltage became positive. There were also isolated periods of small but significant contralateral positivity prior to the negativity.

The multivariate analyses were performed using the same approach used for the N2pc, except that the data were organized according to whether the response was contralateral versus ipsilateral relative to the electrode sites. SVM decoding accuracy was slightly but significantly greater than chance from 324 to 226 ms prior to the response. This was followed by a stronger period of decoding that was significant from 160 ms prior to response until the end of the epoch. A sustained period of significant crossnobis distance began 180 ms prior to the response and lasted through the end of the epoch.

The effect size was similar for SVM decoding accuracy and the univariate voltage during the early part of the LRP (approximately 150-62 ms prior to the response), and then became greater for SVM decoding accuracy until the end of the epoch. The crossnobis effect size was substantially smaller than the univariate effect size prior to the response and then became similar after the response. When the data were averaged across the LRP measurement window (-100 to 0 ms), the effect size was slightly larger for the SVM decoding accuracy than for the univariate voltage and substantially lower for the crossnobis distance (Figure 6C).

### 3.7 ERN

Figure 5D shows the ERN analyses from the flankers paradigm, which examined the difference in voltage between error trials and correct trials. As usual, this analysis yielded a more negative voltage on error trials than on correct trials, which became significant 35 ms before the response and remained significant until 110 ms after the response. The ERN was followed by an error positivity (Pe) that was significant from 132 ms until the end of the epoch. There were also three brief periods of significance prior to the ERN, perhaps reflecting differences in perceptual and cognitive processes that lead to a correct response versus an incorrect response.

The SVM and crossnobis approaches attempted to decode whether the response was correct or incorrect. SVM decoding accuracy and the crossnobis distance became significantly different from chance during this early period prior to the ERN (beginning at 266 ms pre-response for the SVM and at203 ms pre-response for the crossnobis analysis), and they remained significant until the end of the epoch. Note that the univariate analysis made it possible to see separate ERN and Pe components with different polarities, whereas the multivariate analysis did not make this distinction. Prior to the response, the effect size was approximately equal for the two multivariate methods and somewhat smaller for the univariate effect size. After the response, the effect size was greater for the SVM decoding accuracy than for the other two methods. When the data were averaged across the ERN measurement window (0 to 100 ms), the effect size was substantially larger for the SVM decoding accuracy than for the univariate voltage and the crossnobis distance (Figure 6D).

### 3.8 P3b and MMN Results

We chose to describe the results for the P3b and MMN components in the same section because both were assessed using oddball paradigms that are explicitly designed to include one condition with many fewer trials than the other condition. The SVM and crossnobis methods were used to decode whether the stimulus came from the rare category or the frequent category.

Given that our SVM decoding approach requires equating the number of trials across categories by subsampling from the more frequent category, these paradigms may be a special case.

### 3.9 P3b

In the P3b experiment, the univariate data yielded a prototypical P3b effect, consisting of a more positive voltage for the rare targets than for the frequent standards, peaking at approximately 380 ms (see Figure 7A). This effect was significant from 260–660 ms. It was preceded by a much smaller positivity— a frontally-maximal P2 wave—that was significant from 200-226 ms. There was also a brief period of significance from 42-70 ms that is presumably spurious due to its implausibly early onset.

**Figure 7.**
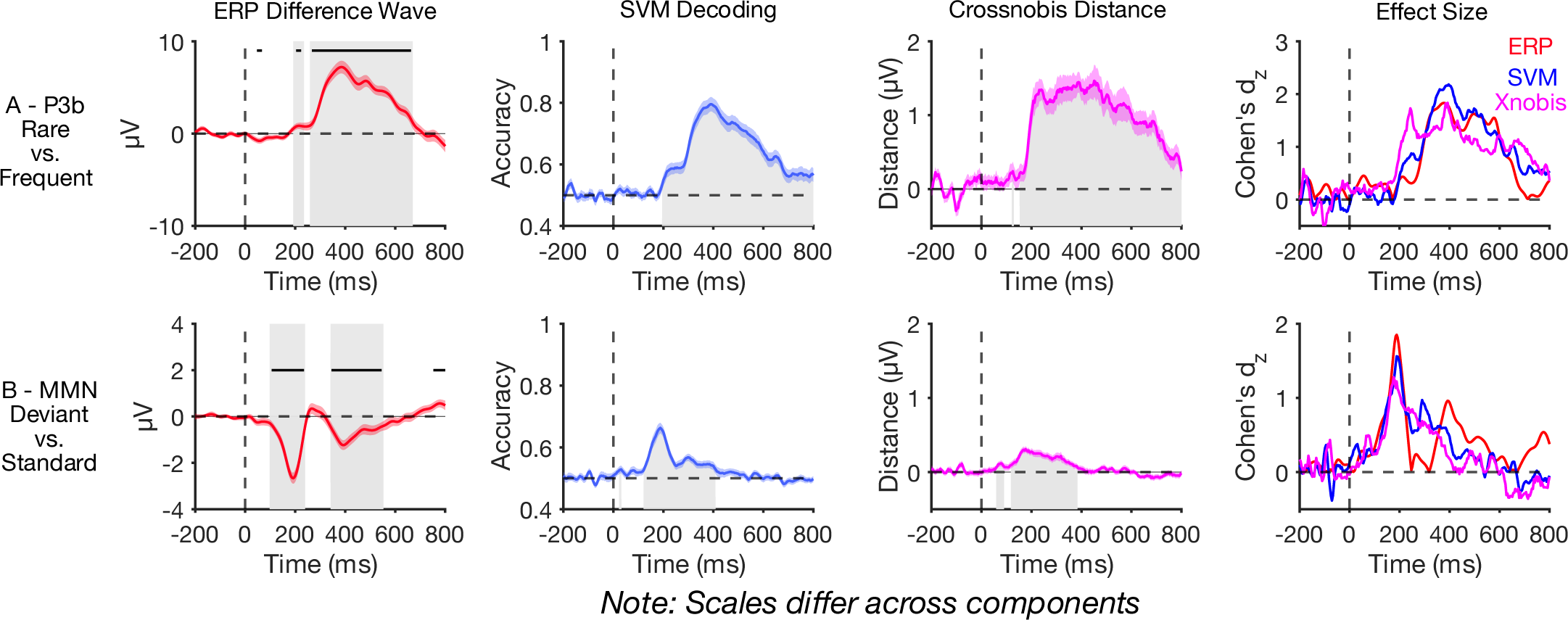
Point-by-point results for the P3b and mismatch negativity (MMN) components. The four columns show the mean ERP difference voltages (averaged across the cluster of electrodes listed in Table 1), mean decoding accuracy values, mean crossnobis distances, and effect sizes at each time point. The shaded bands around the a given waveform show the standard error of the mean. Gray shading indicates time points at which the value is significantly different from chance with a one-tailed test in the direction of the expected effect after correction for multiple comparisons. The horizontal lines for the ERP difference waves indicate time periods at which the value is significantly different from chance with a two-tailed test (which can reveal effects with an unexpected polarity).

The SVM decoding accuracy and crossnobis distance waveforms significantly exceeded chance beginning just before 200 ms, during the time period of the frontal P2 effect, and remained significant through the end of the epoch. During the P2 period, the effect size was much larger for the two multivariate methods than for the univariate voltage, although this may simply reflect the fact that the electrode cluster for the univariate analysis was optimized for the P3b rather than for the more frontally distributed P2. From approximately 300 ms onward, the effect size was similar across methods, but with a small advantage for SVM decoding accuracy near the peak of the P3 wave. When the data were collapsed across the P3b measurement window (300 to 600 ms), the effect size was largest for SVM decoding accuracy, intermediate for the univariate voltage, and smallest for the crossnobis distance (see Figure 8A).

**Figure 8.**
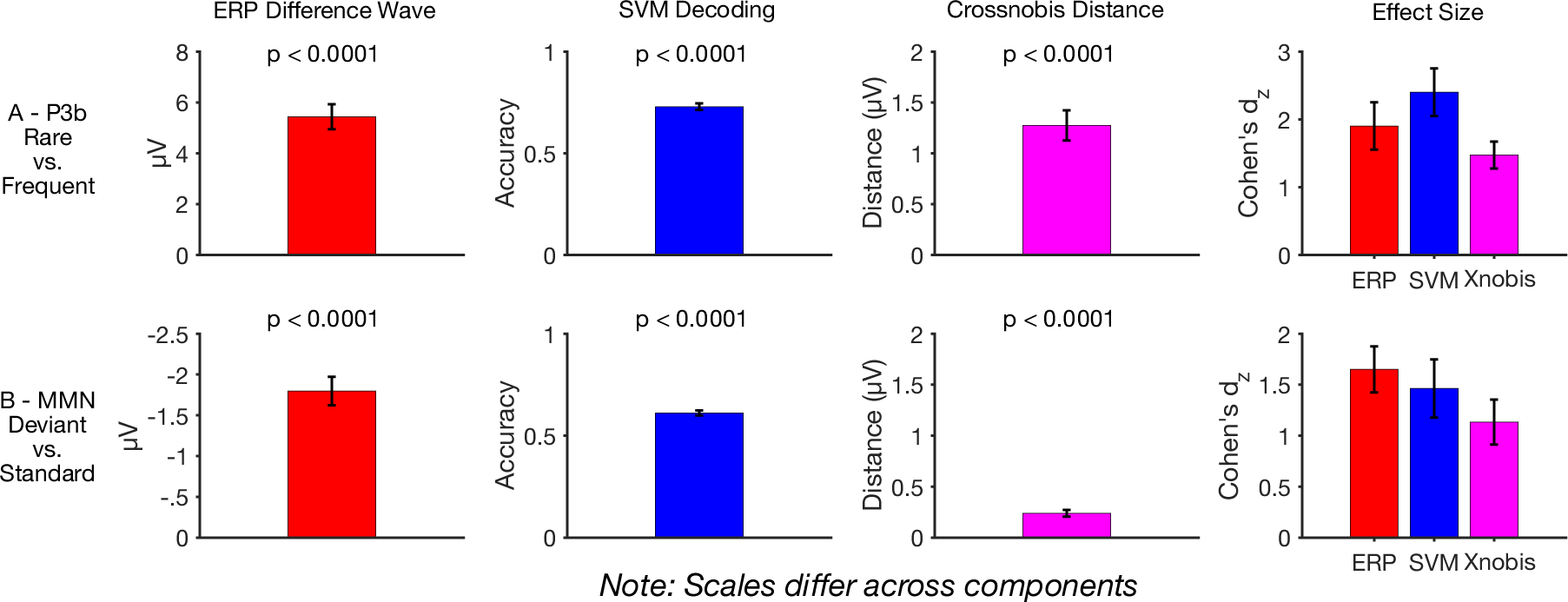
Time-window univariate ERP difference voltage, decoding accuracy, and crossnobis distance, averaged across participants, for the P3b and MMN components. The measurement window was 300–600 ms for the P3b component and 125–225 ms for the MMN. The *p* values are from one-tailed tests against chance. The right column shows the effect size (Cohen’s *d_z_*) at each time window for each analysis method. Error bars show ±1 SEM.

### 3.10 MMN

In the MMN experiment, the univariate data yielded a prototypical MMN effect, consisting of a more negative voltage for deviants than for standards, peaking at approximately 200 ms (see Figure 7B). This effect was significant from 100–240 ms. There was also a significant negativity from 340–546 ms.

The SVM decoding accuracy waveform significantly exceeded chance from 117–406 ms, and the crossnobis distance waveform showed sustained significance from 117–386 ms. The peak effect size was largest for the univariate voltage, slightly smaller for SVM decoding accuracy, and even smaller for crossnobis distance. This same pattern of effect sizes was observed when the data were collapsed across the MMN measurement window (125 to 225 ms; see Figure 8B).

### 3.11 Summary of Effect Sizes and Effect Size Contrasts

Figure 9 summarizes the time-window effect sizes for all the analyses, using the same scale for all components to facilitate comparison across components. The effect size was much larger for the SVM decoding accuracy than for the other approaches in several cases and was never much smaller than the univariate effect size.

**Figure 9.**
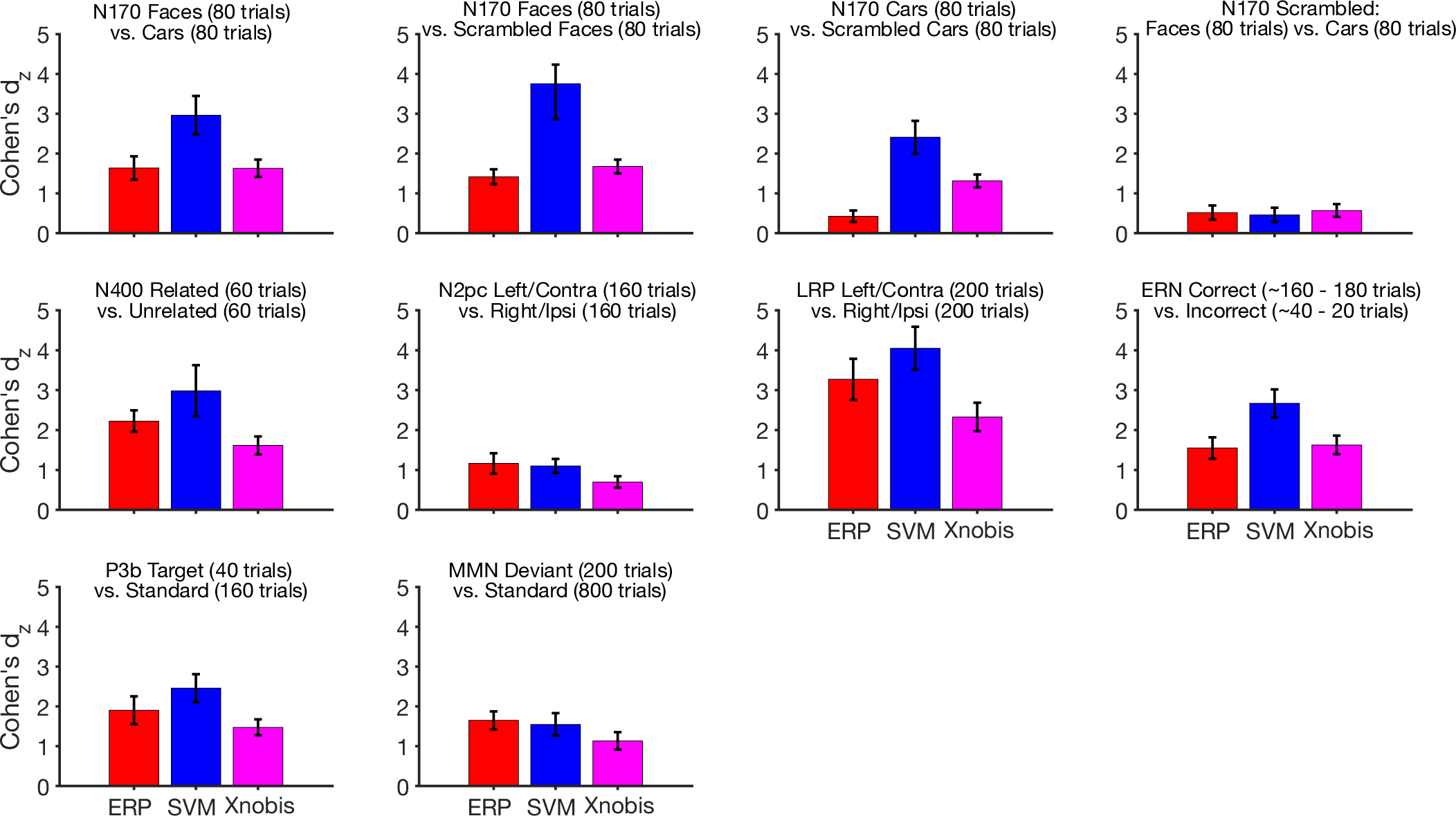
Effect size (Cohen’s *d_z_*) for each analysis method for all the time-window comparisons, drawn on the same scale for all components. The error bars show ±1 standard error.

To provide a direct comparison of the effect sizes, we computed the percentage difference between each pair of analysis methods. The percentage difference was defined as 100 × (*A* – *B*) / *B*, where *A* and *B* represent the effect sizes for two different methods (e.g., SVM decoding accuracy versus univariate difference voltage). We then obtained the 95% confidence intervals of these percentage differences. SVM decoding accuracy yielded the largest effect size in seven of the ten comparisons, figure 10. In these seven cases, the effect size was 14–467% greater for SVM decoding accuracy than for the univariate voltage (with a confidence interval that excluded zero for P3b, ERN, N170 faces versus cars, N170 faces versus scrambled faces, and N170 cars versus scrambled cars). In the other three comparisons, the effect size was only 5%–11% smaller for SVM decoding accuracy than for the univariate voltage, and none of these confidence intervals excluded zero. Thus, SVM decoding accuracy was often much better than the univariate voltage and never much worse, figure 10.

**Figure 10.**
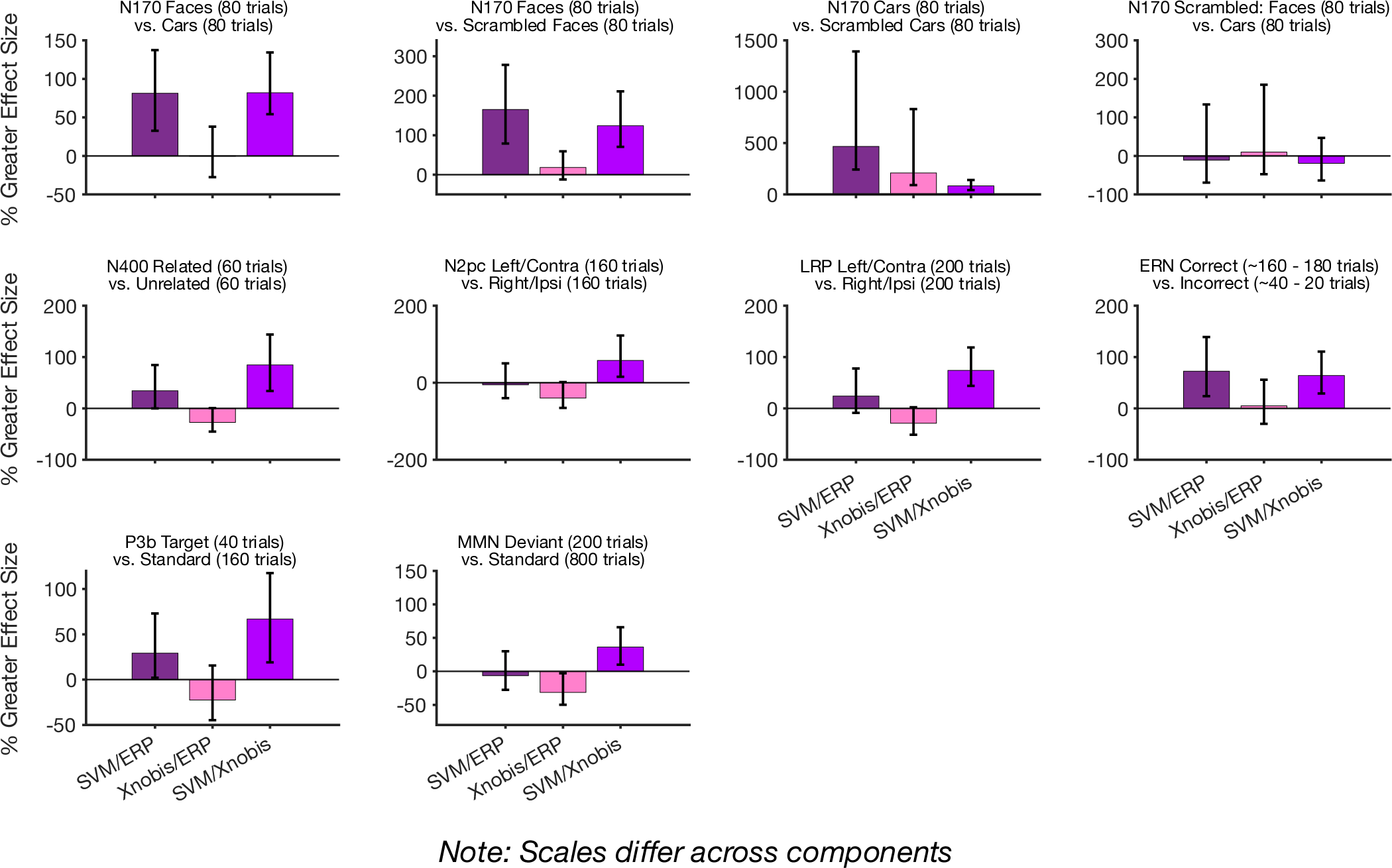
Percentage differences in effect sizes between each pair of analysis methods for all time-window comparisons. The bars show 95% confidence intervals.

The crossnobis distance metric yielded larger effect sizes than the univariate voltage in 4 out of 10 comparisons, for which the benefit was 5–209%. In the remaining 6 comparisons, the effect size was 0.4–40% smaller for the crossnobis distance than for the univariate voltage. Moreover, the SVM effect size was greater than the crossnobis effect size in 9 out of 10 comparisons, with a 36–124% larger effect size for SVM than for crossnobis in these 9 cases. The scrambled faces vs scrambled cars comparison was the only case in which crossnobis yielded a larger effect size than SVM (a 19% advantage, with a confidence interval that included zero), figure 10.

### 3.12 Mean Amplitude × Multivariate Time Window Regressions

The univariate and multivariate analysis methods were applied to the same data, and they assessed the same or similar contrasts between experimental conditions, so it seems plausible that participants who have a large univariate difference in voltage between conditions will also exhibit high SVM decoding accuracy and large crossnobis distances. However, the SVM and crossnobis methods also take into account the variability across trials within each participant, so a participant with a large difference in mean voltage between two conditions but high trial-to-trial variability might have low decoding accuracy and a small crossnobis distance. Thus, it is an empirical question whether the size of the univariate voltage difference will be associated with the SVM decoding accuracy or the crossnobis distance. We therefore used simple linear regressions to assess the strength of these associations. The polarities of the plots and the *r* values were reversed when necessary, so that a larger univariate effect was always coded as a more positive value.

As shown in Figure 11A, the univariate voltage difference between conditions was significantly correlated with the SVM decoding accuracy across all ten comparisons. The magnitude of the correlations varied widely from largely medium to strong correlations (from *r* = 0.344 to 0.791), but the correlation was above 0.4 for all components except the N170 faces vs scrambled faces comparison which is where the univariate effect size was quite small.

**Figure 11.**
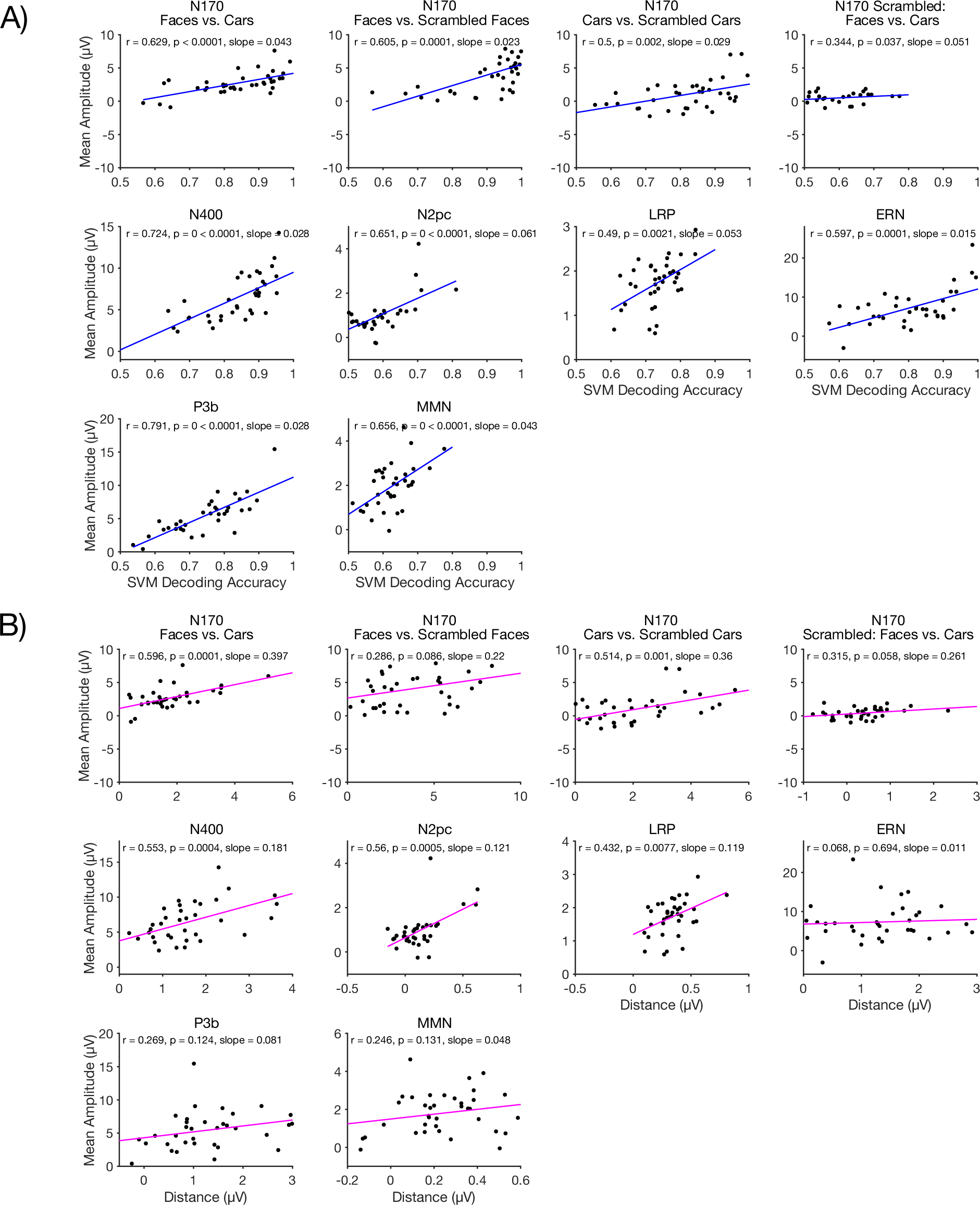
Difference wave mean amplitude as a function of SVM decoding accuracy (A) or crossnobis distance (B). Note that the polarities of the plots and *r* values were reversed when necessary, so that a larger univariate effect was always coded as a more positive value. The Pearson *r* correlation coefficient, corresponding *p* value, and slope of the linear relationship are shown for each comparison.

Figure 11B shows the correlations between the univariate voltage differences and the crossnobis distances. These correlations were much smaller than those for the SVM decoding accuracy, and they failed to reach significance in six of the ten cases (N170 faces vs. scrambled faces; N170 scrambled faces vs. scrambled cars; ERN, P3b, and MMN).

## 4. General Discussion

In this study, we compared the traditional univariate approach to assessing differences in amplitude between experimental conditions to newer multivariate pattern analysis methods that apply machine learning to the entire topographic pattern of voltage across the scalp. We examined ten different comparisons across seven different ERP components so that we could assess the generality of any differences. When the data were averaged across the measurement window for the component of interest, we found that SVM decoding accuracy produced substantially larger effect sizes than traditional univariate methods in seven of the ten comparisons, with a slightly smaller effect size for SVM decoding accuracy in the other three cases. Similar results were obtained in point-by-point analyses, although there was some variation in the relative effect sizes across time points. In almost every case, effect sizes were lower for the crossnobis distance than for SVM decoding accuracy. These results suggest that ERP researchers could obtain larger effect sizes, and therefore greater statistical power, by using SVM decoding accuracy as their primary dependent variable instead of the univariate voltage, at least for the types of participants and paradigms examined in the present study. Moreover, SVM decoding is now very straightforward to apply in ERPLAB Toolbox (Lopez-Calderon & Luck, 2014), using either scripting or a point-and-click graphical user interface.

Larger effect sizes and greater statistical power would be a substantial boon to ERP research. A larger effect size can decrease the Type II error rate and thereby increase the proportion of significant effects that are actually true effects (Ioannidis, 2005). Alternatively, researchers could achieve the same level of statistical power as before with fewer participants, which would increase the number of studies that are feasible given limited resources. However, there are some important limitations to the present findings and to the use of SVM decoding in place of traditional univariate analyses, so researchers should carefully consider whether to replace conventional analyses with SVM decoding. We will focus the remainder of our discussion on these limitations.

Although the present study used many different comparisons and ERP components to enhance the generality of our conclusions, the benefits of SVM decoding we observed may not generalize to all ERP research. The ERP CORE data used for the present analyses were recorded using active electrodes with a stable gel-based interface between the electrodes, and the recordings were obtained inside an electrically shielded chamber (which minimizes line noise) with excellent temperature control (which minimizes skin potentials). Moreover, the data were obtained from highly compliant neurotypical young adults. Because SVM decoding accuracy and the crossnobis distance are both decreased when trial-to-trial variability in the EEG is greater, they may yield smaller effect sizes than univariate voltage measures in datasets with greater noise. However, univariate effect sizes are also decreased when the trial-to-trial variability increases. It would therefore be useful for future research to conduct similar comparisons between univariate and multivariate methods in data obtained in noisier recording setups and in data from participant populations who exhibit greater trial-to-trial EEG variability.

A second limit to generality is that we cannot be certain that SVM decoding accuracy would lead to larger effect sizes for all ERP paradigms. Indeed, we found that it led to comparable or slightly smaller effect sizes than the univariate voltage in a few cases. It is not clear why. The fact that SVM decoding was not very helpful for the N2pc and LRP components may reflect the fact that the traditional univariate approach already uses a difference in scalp distribution (contralateral versus ipsilateral) to quantify the amplitude of the component. However, this does not explain why SVM decoding had no advantage over the univariate voltage for the MMN or for the comparison of scrambled faces and scrambled cars in the N170 experiment. These variations in the relative effectiveness of SVM decoding may reflect some as-yet-to-be-determined variable or they may reflect random variation. It will be important for future research to determine whether there are specific conditions under which SVM decoding is advantageous. However, SVM decoding was never substantially worse than univariate voltage analyses in the present data, and it was often substantially better, so researchers who are analyzing similar data should seriously consider using SVM decoding.

A key difference between traditional univariate analyses and MVPA analyses is that univariate analyses often focus on the subset of electrodes where the effect is largest, whereas MVPA analyses typically use all electrodes. Focusing on electrodes with large effects might be expected to confer an advantage, but we found that including all the sites in the MVPA analyses led to equivalent or even larger effect sizes than including only the sites from the univariate analyses (see supplementary Figures S5 – S10).

Another increasingly common univariate approach is the mass univariate approach, in which conditions are statistically compared separately at each electrode site, and then an intelligent correction for multiple comparisons is applied (Maris & Oostenveld, 2007). There is no straightforward way to assess the effect size with this approach, which makes it difficult to compare with MVPA approaches. However, the statistical power of a mass univariate analysis is typically lower than the power of the univariate approach used in the present study, because confining a univariate analysis to electrode sites that are known a priori to contain the largest effects increases the power of the analysis relative to a mass univariate analysis (Groppe et al., 2011). Thus, we would expect that SVM-based decoding analyses would have even greater advantages over mass univariate analyses than over the simple univariate analyses examined here.

A related issue is that univariate analyses provide scalp distribution information, whereas SVM-based decoding does not. For example, studies using univariate approaches often include scalp maps of the effects (including all electrodes, not just those used in the analyses). Moreover, the mass univariate approach can indicate which sites exhibit statistically significant effects. This information is not provided by SVM-based decoding. Although it is possible to look at the weight for each electrode site in an SVM analysis, Haufe et al. (2014) showed that this can be misleading. They proposed using a transformation of the weights, but this transformation applies only to linear discriminant analysis, not to SVM-based decoding. In addition, MVPA approaches are applied separately to each participant, so there is no group-level scalp map. However, this single-participant approach is also an advantage, because it is not hampered by differences across participants in the scalp distribution of an effect, which may be quite large due to idiosyncrasies in biophysical factors (Hajizadeh et al., 2021). This may be a key reason why SVM-based decoding often led to larger effect sizes than univariate methods in the present study.

The fact that SVM decoding requires equating the number of trials between the conditions being compared is a potential downside and may explain the relatively poor performance of SVM decoding for the MMN (see Luck, 2023 for an extended discussion). However, we obtained larger effect sizes for SVM decoding than for the univariate voltage for the P3b and ERN components, which also required discarding large numbers of trials in one of the two conditions being compared. It should be noted that care would be needed in applying SVM decoding in studies comparing groups of participants that differ in the number of trials available for averaging (e.g., due to differences in the number of trials rejected because of artifacts). In such studies, it would be necessary to match the number of trials per groups by subsampling from the available trials. This could be accomplished by pairing participants across groups and using subsampling to equate the number of trials in each pair.

Increased trial-to-trial variability does not impact the expected value of traditional mean amplitude measures (Luck, 2014). That is, although the measured mean amplitude may be less representative of the true value in noisier data, it is not consistently larger or consistently smaller in noisier data than in cleaner data. However, the expected value for SVM decoding accuracy becomes progressively smaller as the trial-to-trial variability increases. As a result, it can be problematic to compare decoding accuracy across groups or experimental conditions that differ in the amount of trial-to-trial variability. For example, it may be problematic to compare decoding accuracy between a patient group and a control group if the patient group has more movement artifacts than the control group. A method has been developed to quantify the specific type of trial-to-trial variability that impacts SVM decoding accuracy (Bae et al., 2023), and a method of this nature should be used whenever there is reason to believe that groups or conditions might differ in their variability. However, this is not a problem if decoding accuracy is actually greater in the group with greater variability than the group with less variability. It should also be noted that this issue is not unique to decoding accuracy: The expected value of traditional peak amplitude measures increases progressively as the noise level increases, making it problematic to compare groups with different noise levels (unless the group with a higher noise level is found to have lower peak amplitudes than the group with a lower noise level).

Another important limitation of decoding is the loss of polarity information. In the case of the N2pc component, for example, a contralateral positivity often follows the N2pc and is thought to reflect a different cognitive process than the N2pc (Gaspelin et al., 2023). However, the SVM and crossnobis analyses are blind to polarity and simply indicate that the two conditions are discriminable across a set of time points.

On the other hand, it can be very difficult to isolate a specific ERP component and to know whether the same component is being observed across different experimental paradigms and participant populations (Kappenman & Luck, 2012b). Decoding approaches avoid these thorny issues, instead asking whether two experimental conditions yield discriminably different patterns of neural activity, how different they are, and the latencies at which they are discriminable.

Indeed, the interpretation of decoding accuracy is subtly but importantly different than the traditional interpretation of univariate voltage differences. Univariate ERP analyses are traditionally used to quantify the extent to which *a specific ERP component* differs across conditions, with the implicit assumption that the component has a relatively constant scalp distribution across participants (and can therefore be measured at the same electrode or cluster of electrodes in all participants). Decoding analyses instead quantify *the amount of information* about condition that is present in the recorded signal in each individual participant, with no assumption that the underlying brain activity is similar across participants. These are both valid things to quantify, and which one is more useful will depend on the scientific question being asked by a given study. Researchers must therefore think carefully about whether their scientific question requires isolating a specific component or whether it is instead related to the information content of the neural signal.

## Data and Code Availability

All raw and preprocessed univariate data used for this project are a part of the ERP CORE resource, which is available at https://doi.org/10.18115/D5JW4R. The data and code used to implement all decoding analyses that is compatible with ERPLAB’s GUI is available here: doi.org/10.17605/OSF.IO/5WV93.

## Author Contributions

Carlos Daniel Carrasco: conceptualization, analyses, writing; Brett Bahle: development of crossnobis method, crossnobis description writing, Aaron Matthew Simmons: software development; Steven J. Luck: conceptualization, writing, development of methods, funding acquisition. Editing/Review was performed by all authors prior to submission of the final version of the manuscript.

## Acknowledgements

We thank John Nadra for always providing feedback and support throughout the duration of this project and many others. We also thank the members of the Luck Lab for their valuable contributions to community and thoughtful insights, including but not limited to: David Garrett, Guanghui Zhang, Lara Krisst, Kurt Winsler, and John Kiat.

## Footnotes

1 Here, we focus on different classes of stimuli, but these methods can also be applied to classes based on other factors, such as responses (e.g., left-hand vs. right-hand responses). The only restriction is that the methods discussed here require discrete classes. Regression-based methods can be used to decode continuous variables (e.g., Collins & Frank, 2018).

2 It is possible to apply a dimensionality reduction method such as principal component analysis to the data prior to decoding, but we have not found much advantage to this. It is also possible to use a nonlinear hyperplane to distinguish between the classes. However, when an SVM is applied to ERP data in which each dimension represents the same type of variable (voltage) and a large number of dimensions (electrodes) is used, we find that a linear hyperplane typically works best.

3 In this paper, we will focus on binary classifications in which there are two classes being distinguished (e.g., faces versus cars). However, it is straightforward to extend the approaches discussed here to multiclass problems in which three or more classes are being distinguished (e.g., which of 10 faces is being perceived). This can be achieved by decoding each possible pair and averaging the accuracy across all pairs. However, we have found better results using an approach called *error-correcting output codes* (Dietterich & Bakiri, 1995), in which a set of decoders is combined to achieve a single decision for each test case (Bae & Luck, 2018; Bae, 2019; Bae & Luck, 2021).

4 The location of the reference site(s) has little or no impact on the decoding methods used here

5 This subsampling procedure is also widely used in conventional univariate analyses comparing peak amplitudes, because peak values are biased to be larger when the waveform is noisier. However, subsampling is not necessary for mean amplitudes, which are not consistently larger for noisier data (Luck, 2014). Our goal in the present study was to compare the standard univariate approach (which does not require subsampling for mean amplitude) with the standard SVM approach (which does require subsampling).

6 In the univariate analysis, the voltage at a given point in the difference wave might be significantly greater than zero or significantly less than zero, so a two-tailed test was necessary. For decoding, either a positive or a negative difference would yield an above-chance decoding accuracy value, so a one-tailed test was appropriate. We are not comparing significance values between the univariate and decoding methods, so this does not give decoding an unfair advantage.

7 A consequence of this is that the value inside the square root operation can be either positive or negative. To avoid the use of imaginary numbers, the absolute value is taken prior to computing the square root. The original sign of the value inside the square root operation is then applied to the result.

## References

Ashton, K., Zinszer, B. D., Cichy, R. M., Nelson, C. A., Aslin, R. N., & Bayet, L. (2022). Time-resolved multivariate pattern analysis of infant EEG data: A practical tutorial. Developmental Cognitive Neuroscience, 54. 10.1016/j.dcn.2022.101094

Bae, G-Y, Leonard, C. J., Hahn, B., Gold, J. M., & Luck, S. J. (2023). Assessing the information content of ERP signals in schizophrenia using multivariate decoding methods. NeuroImage: Clinical, 25. 10.1016/j.nicl.2020.102179

Bae, G-Y, & Luck, S. J. (2019). Decoding motion direction using the topography of sustained ERPs and alpha oscillations. NeuroImage, 184, 242–255. 10.1016/j.neuroimage.2018.09.029

Bae, G-Y (2021). The Time Course of Face Representations during Perception and Working Memory Maintenance. Cerebral Cortex Communications, 2(1), 1–12. 10.1093/texcom/tgaa093

Bae, G-Y, & Luck, S. J. (2018). Dissociable Decoding of Spatial Attention and Working Memory from EEG Oscillations and Sustained Potentials. The Journal of Neuroscience, 38(2), 2860–17. 10.1523/JNEUROSCI.2860-17.2017

Benjamini, Y., & Yekutieli, D. (2001). The control of the false discovery rate in multiple testing under dependency. The Annals of Statistics, 29(4), 1165–1188. 10.1214/aos/1013699998

Borrell, V. (2018). How cells fold the cerebral cortex. Journal of Neuroscience, 38(4), 776–783. 10.1523/JNEUROSCI.1106-17.2017

Canty, A., & Ripley, B. D. (2022). boot: Bootstrap R (S-Plus) Functions.

Chan, A. M., Halgren, E., Marinkovic, K., & Cash, S. S. (2011). Decoding word and category-specific spatiotemporal representations from MEG and EEG. NeuroImage, 54(4), 3028– 3039. 10.1016/j.neuroimage.2010.10.073

Cohen J. (1988). Statistical Power Analysis for the Behavioral Sciences Second Edition.

Collins, A. G. E., & Frank, M. J. (2018). Within- and across-trial dynamics of human EEG reveal cooperative interplay between reinforcement learning and working memory. Proceedings of the National Academy of Sciences of the United States of America, 115(10), 2502–2507. 10.1073/pnas.1720963115

Daly, I. (2023). Neural decoding of music from the EEG. Scientific Reports, 13(1). 10.1038/s41598-022-27361-x

Davison, A. C., & Hinkley, D. V. (1997). Bootstrap Methods and Their Applications. Cambridge University Press. http://statwww.epfl.ch/davison/BMA/

de Vries, I. E. J., van Driel, J., & Olivers, C. N. L. (2019). Decoding the status of working memory representations in preparation of visual selection. NeuroImage, 191, 549–559. 10.1016/j.neuroimage.2019.02.069

Delorme, A., & Makeig, S. (2004). EEGLAB: an open source toolbox for analysis of single-trial EEG dynamics including independent component analysis. In Journal of Neuroscience Methods (Vol. 134). http://www.sccn.ucsd.edu/eeglab/

Diciccio, T. J., & Efron, B. (1996). Bootstrap Confidence Intervals. In Statistical Science (Vol. 11, Issue 3).

Dietterich, T. G., & Bakiri, G. (1995). Solving Multiclass Learning Problems via Error-Correcting Output Codes. In Journal of Artiicial Intelligence Research (Vol. 2).

Dobs, K., Isik, L., Pantazis, D., & Kanwisher, N. (2019). How face perception unfolds over time. Nature Communications, 10(1). 10.1038/s41467-019-09239-1

Efron, B., & Tibshirani, R. J. (1994). An introduction to the bootstrap. CRC press.

Eimer, M., & Coles, M. G. H. (2003). The lateralized readiness potential. The Bereitschaftspotential: Movement-Related Cortical Potentials, 229–248.

Ester, E., & Weese, R. (2023). Temporally Dissociable Mechanisms of Spatial, Feature, and Motor Selection during Working Memory-guided Behavior. Journal of Cognitive Neuroscience, 1–14. 10.1162/jocn_a_02061

Fernández, V., Llinares-Benadero, C., & Borrell, V. (2016). Cerebral cortex expansion and folding: what have we learned? The EMBO Journal, 35(10), 1021–1044. 10.15252/embj.201593701

Gaspelin, N., Lamy, D., Egeth, H. E., Liesefeld, H. R., Kerzel, D., Mandal, A., Müller, M. M., Schall, J. D., Schubö, A., Slagter, H. A., Stilwell, B. T., & van Moorselaar, D. (2023). The Distractor Positivity Component and the Inhibition of Distracting Stimuli. Journal of Cognitive Neuroscience, 1–23. 10.1162/jocn_a_02051

Gehring, W. J., Liu, Y., Orr, J. M., & Carp, J. (2012). The error-related negativity (ERN/Ne). Oxford Handbook of Event-Related Potential Components, 231–291.

Gholami, R., & Fakhari, N. (2017). Support Vector Machine: Principles, Parameters, and Applications. In Handbook of Neural Computation (pp. 515–535). Elsevier Inc. 10.1016/B978-0-12-811318-9.00027-2

Grootswagers, T., Wardle, S. G., & Carlson, T. A. (2017). Decoding dynamic brain patterns from evoked responses: A tutorial on multivariate pattern analysis applied to time series neuroimaging data. Journal of Cognitive Neuroscience, 29(4), 677–697. 10.1162/jocn_a_01068

Groppe, D. M., Urbach, T. P., & Kutas, M. (2011). Mass univariate analysis of event-related brain potentials/fields I: A critical tutorial review. In Psychophysiology (Vol. 48, Issue 12, pp. 1711–1725). Blackwell Publishing Inc. 10.1111/j.1469-8986.2011.01273.x

Gurariy, G., Mruczek, R. E. B., Snow, J. C., & Caplovitz, G. P. (2022). Using High-Density Electroencephalography to Explore Spatiotemporal Representations of Object Categories in Visual Cortex. Journal of Cognitive Neuroscience, 34(6), 967–987. 10.1162/jocn_a_01845

Hajizadeh, A., Matysiak, A., Brechmann, A., König, R., & May, P. J. C. (2021). Why do humans have unique auditory event-related fields? Evidence from computational modeling and MEG experiments. Psychophysiology, 58(4). 10.1111/psyp.13769

Haufe, S., Meinecke, F., Görgen, K., Dähne, S., Haynes, J. D., Blankertz, B., & Bießmann, F. (2014). On the interpretation of weight vectors of linear models in multivariate neuroimaging. NeuroImage, 87, 96–110. 10.1016/j.neuroimage.2013.10.067

Hubbard, J., Kikumoto, A., & Mayr, U. (2019). EEG Decoding Reveals the Strength and Temporal Dynamics of Goal-Relevant Representations. Scientific Reports, 9(1). 10.1038/s41598-019-45333-6

Ioannidis, J. P. A. (2005). Why most published research findings are false. In Why most published research findings are false. (Vol. e124, pp. 2–8). Springer International Publishing. 10.1371/journal.pmed.0020124

Kappenman, E. S., Farrens, J. L., Zhang, W., Stewart, A. X., & Luck, S. J. (2021). ERP CORE: An open resource for human event-related potential research. NeuroImage, 225(May 2020), 117465. 10.1016/j.neuroimage.2020.117465

Kappenman, E. S., & Luck, S. J. (2012a). Electrophysiological correlates of the focusing of attention within complex visual scenes: N2pc and related ERP components. In The Oxford Handbook of Event-Related Potential Components (pp. 329–360). Oxford University Press. 10.1093/oxfordhb/9780195374148.001.0001

Kappenman, E. S., & Luck, S. J. (2012b). ERP components: The ups and downs of brainwave recordings. In The Oxford handbook of event-related potential components. (pp. 3–30). Oxford University Press.

Kiss, M., Van Velzen, J., & Eimer, M. (2008). The N2pc component and its links to attention shifts and spatially selective visual processing. Psychophysiology, 45(2), 240–249. 10.1111/j.1469-8986.2007.00611.x

Kroenke, C. D., & Bayly, P. V. (2018). How forces fold the cerebral cortex. Journal of Neuroscience, 38(4), 767–775. 10.1523/JNEUROSCI.1105-17.2017

Kutas, M., & Federmeier, K. D. (2011). Thirty years and counting: Finding meaning in the N400 component of the event-related brain potential (ERP). Annual Review of Psychology, 62, 621–647. 10.1146/annurev.psych.093008.131123

Lakens, D. (2013). Calculating and reporting effect sizes to facilitate cumulative science: A practical primer for t-tests and ANOVAs. Frontiers in Psychology, 4(NOV). 10.3389/fpsyg.2013.00863

Li, D., Hu, Y., Qi, M., Zhao, C., Jensen, O., Huang, J., & Song, Y. (2023). Prioritizing flexible working memory representations through retrospective attentional strengthening. NeuroImage, 269. 10.1016/j.neuroimage.2023.119902

Li, Y., Zhang, M., Liu, S., & Luo, W. (2022). EEG decoding of multidimensional information from emotional faces. NeuroImage, 258. 10.1016/j.neuroimage.2022.119374

Lopez-Calderon, J., & Luck, S. J. (2014). ERPLAB: An open-source toolbox for the analysis of event-related potentials. Frontiers in Human Neuroscience, *8*(1 APR), 1–14. 10.3389/fnhum.2014.00213

Luck, S. J. (2014). An Introduction To The Event-Related Potential Technique (2nd Editio). MIT Press Journals.

Luck, S. J. (2023). ERPLAB Decoding Tutorial. https://github.com/ucdavis/erplab/wiki/ERPLAB-Decoding-Tutorial

Maris, E., & Oostenveld, R. (2007). Nonparametric statistical testing of EEG-and MEG-data. Journal of Neuroscience Methods, 164(1), 177–190. 10.1016/j.jneumeth.2007.03.024

MathWorks. (2023). MATLAB version: 23.2.0 (R2023b). The MathWorks Inc. https://www.mathworks.com

Mazza, V., Turatto, M., Umiltà, C., & Eimer, M. (2007). Attentional selection and identification of visual objects are reflected by distinct electrophysiological responses. Experimental Brain Research, 181(3), 531–536. 10.1007/s00221-007-1002-4

Murphy, B., Poesio, M., Bovolo, F., Bruzzone, L., Dalponte, M., & Lakany, H. (2011). EEG decoding of semantic category reveals distributed representations for single concepts. Brain and Language, 117(1), 12–22. 10.1016/j.bandl.2010.09.013

Näätänen, R., Paavilainen, P., Rinne, T., & Alho, K. (2007). The mismatch negativity (MMN) in basic research of central auditory processing: A review. In Clinical Neurophysiology (Vol. 118, Issue 12, pp. 2544–2590). 10.1016/j.clinph.2007.04.026

Nadra, J. G., Bengson, J. J., Morales, A. B., & Mangun, G. R. (2023). Attention without Constraint: Alpha Lateralization in Uncued Willed Attention. ENeuro, 10(6). 10.1523/ENEURO.0258-22.2023

Nemrodov, D., Niemeier, M., Patel, A., & Nestor, A. (2018). The neural dynamics of facial identity processing: Insights from EEG-based pattern analysis and image reconstruction. ENeuro, 5(1). 10.1523/ENEURO.0358-17.2018

Noah, S., Powell, T., Khodayari, N., Olivan, D., Ding, M., & Mangun, G. R. (2020). Neural Mechanisms of Attentional Control for Objects: Decoding EEG Alpha When Anticipating Faces, Scenes, and Tools. Journal of Neuroscience, 40(25), 4913–4924. 10.1523/JNEUROSCI.2685-19.2020

Polich, J. (2012). Neuropsychology of P300. The Oxford Handbook of Event-Related Potential Components, 159–188.

Prime, D. J., & Jolicoeur, P. (2010). Mental Rotation Requires Visual Short-term Memory: Evidence from Human Electric Cortical Activity. Journal of Cognitive Neuroscience. 10.1162/jocn.2009.21337

R Core Team. (2023). R: A Language and Environment for Statistical Computing. https://www.R-project.org/

Rossion, B., & Jacques, C. (2011). The N170: Understanding the time course of face perception in the human brain. The Oxford Handbook of ERP Components, 115–142.

Sawaki, R., Geng, J. J., & Luck, S. J. (2012). A common neural mechanism for preventing and terminating the allocation of attention. Journal of Neuroscience, 32(31), 10725–10736. 10.1523/JNEUROSCI.1864-12.2012

Smith, F. W., & Smith, M. L. (2019). Decoding the dynamic representation of facial expressions of emotion in explicit and incidental tasks. NeuroImage, 195, 261–271. 10.1016/j.neuroimage.2019.03.065

Trammel, T., Khodayari, N., Luck, S. J., Traxler, M. J., & Swaab, T. Y. (2023). Decoding semantic relatedness and prediction from EEG: A classification method comparison. NeuroImage, 277, 120268. 10.1016/j.neuroimage.2023.120268

Walther, A., Nili, H., Ejaz, N., Alink, A., Kriegeskorte, N., & Diedrichsen, J. (2016). Reliability of dissimilarity measures for multi-voxel pattern analysis. NeuroImage, 137, 188–200. 10.1016/j.neuroimage.2015.12.012

Wang, D., Miao, D., & Blohm, G. (2012). Multi-class motor imagery EEG decoding for brain-computer interfaces. *Frontiers in Neuroscience*, OCT. 10.3389/fnins.2012.00151

Wolff, M. J., Ding, J., Myers, N. E., & Stokes, M. G. (2015). Revealing hidden states in visual working memory using electroencephalography. Frontiers in Systems Neuroscience, 9(september). 10.3389/fnsys.2015.00123

Yang, L., Song, Y., Ma, K., & Xie, L. (2021). Motor Imagery EEG Decoding Method Based on a Discriminative Feature Learning Strategy. IEEE Transactions on Neural Systems and Rehabilitation Engineering, 29, 368–379. 10.1109/TNSRE.2021.3051958

Zhang, W., & Kappenman, E. S. (2023). Maximizing signal-to-noise ratio and statistical power in ERP measurement: Single sites versus multi-site average clusters. Psychophysiology. 10.1111/psyp.14440

