## Supplemental for "Using Multivariate Pattern Analysis to Increase Effect Sizes for Event-Related Potential Analyses"

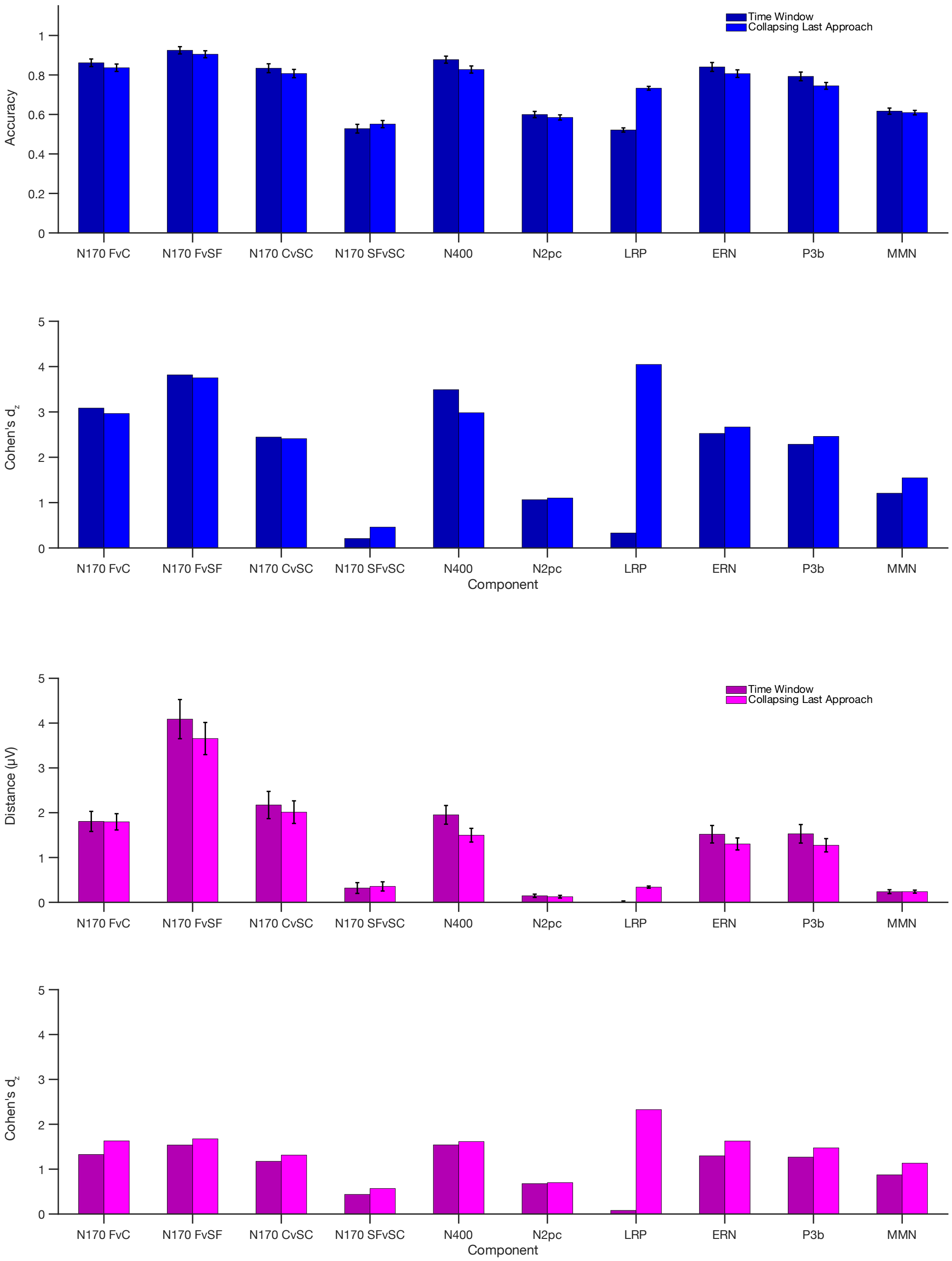


Figure S1. Comparison of two approaches for computing time-window values for the SVM and crossnobis methods. In the collapsing-last approach, we simply performed decoding on the individual time points and averaged the point-by-point decoding accuracy values and crossnobis distances across the measurement window. In the collapsing-first approach, we averaged the voltages across the time window in each single-trial epoch prior to the decoding and crossnobis analyses.

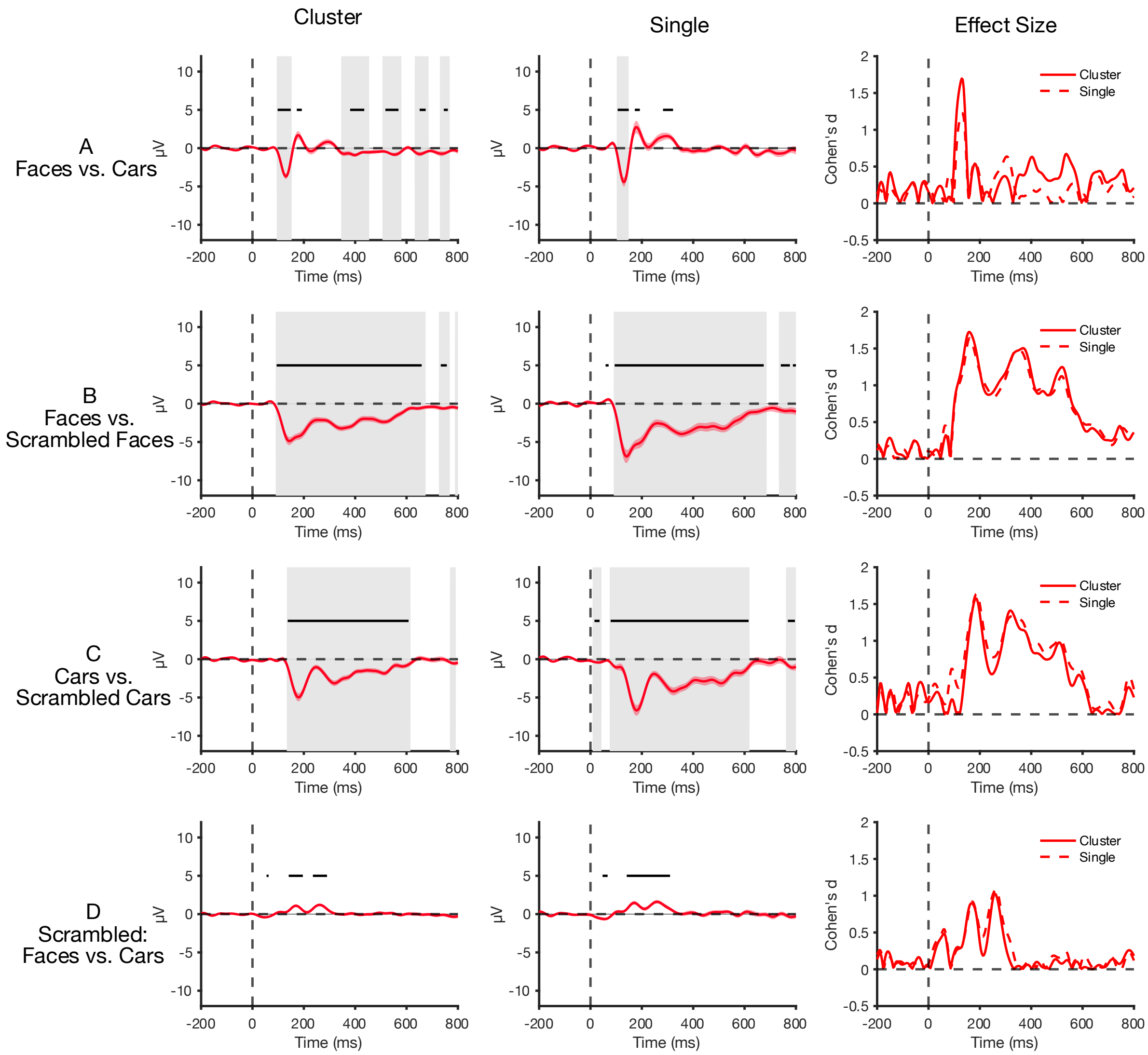


Figure S2. Point-by-point ERP difference voltages, for the 4 components elicited by the N170 paradigm using single sites, and clusters of sites. The shaded bands around the waveform show the standard error of the mean. Gray shading indicates significance at $p < 0.05$, FDR corrected. The right column shows the effect size (Cohen's $d_z$) at each time point for each analysis method. The cluster sites tend to yield larger effect sizes than using single sites.

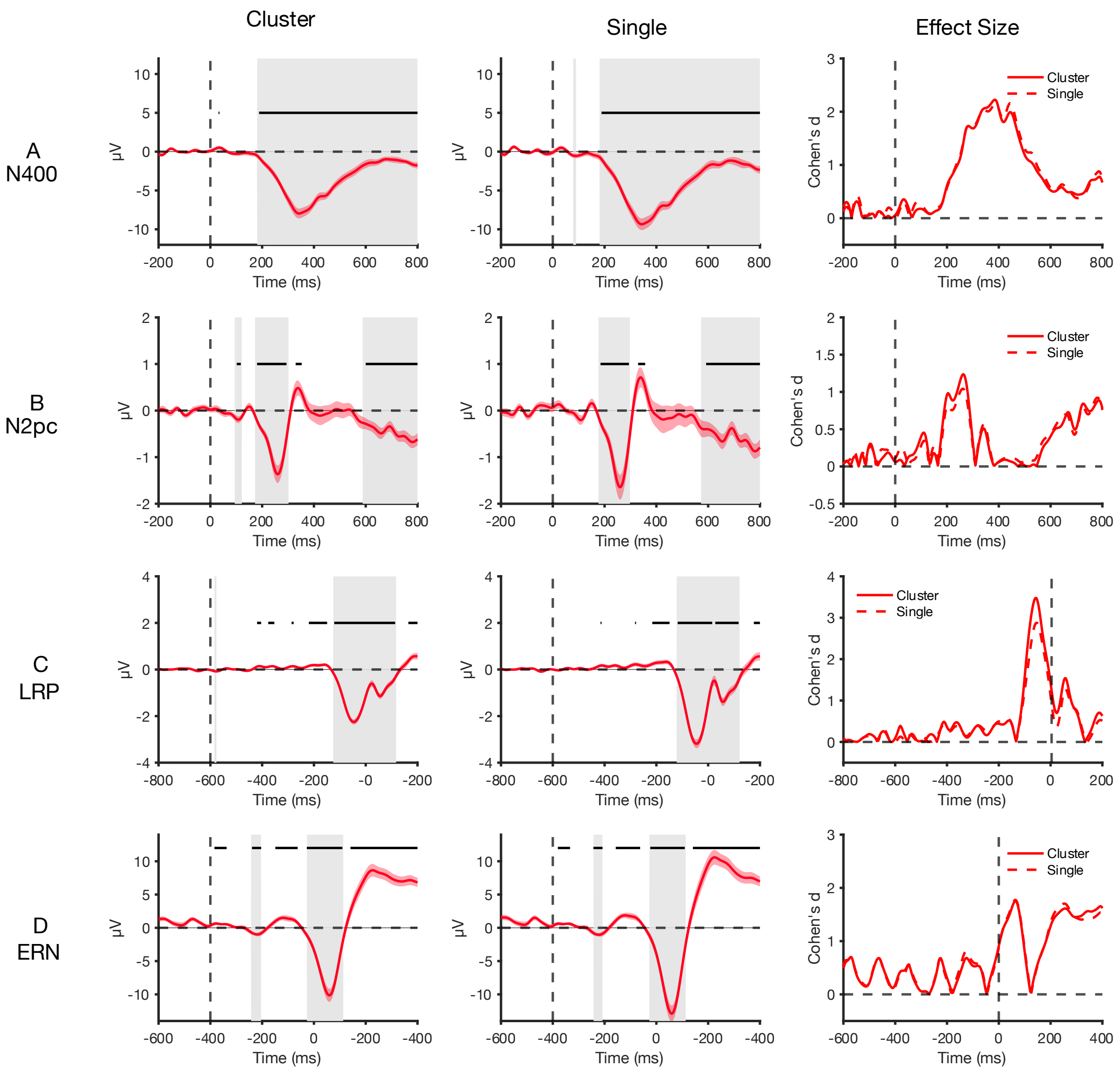


Figure S3. Point-by-point ERP difference voltages, for the 4 components elicited by the N400, N2pc, LRP/ERN paradigms using single sites and clusters of sites. The shaded bands around the waveform show the standard error of the mean. Gray shading indicates significance at $p < 0.05$, FDR corrected. The right column shows the effect size (Cohen's $d_z$) at each time point for each analysis method. The cluster sites tend to yield larger effect sizes than using single sites.

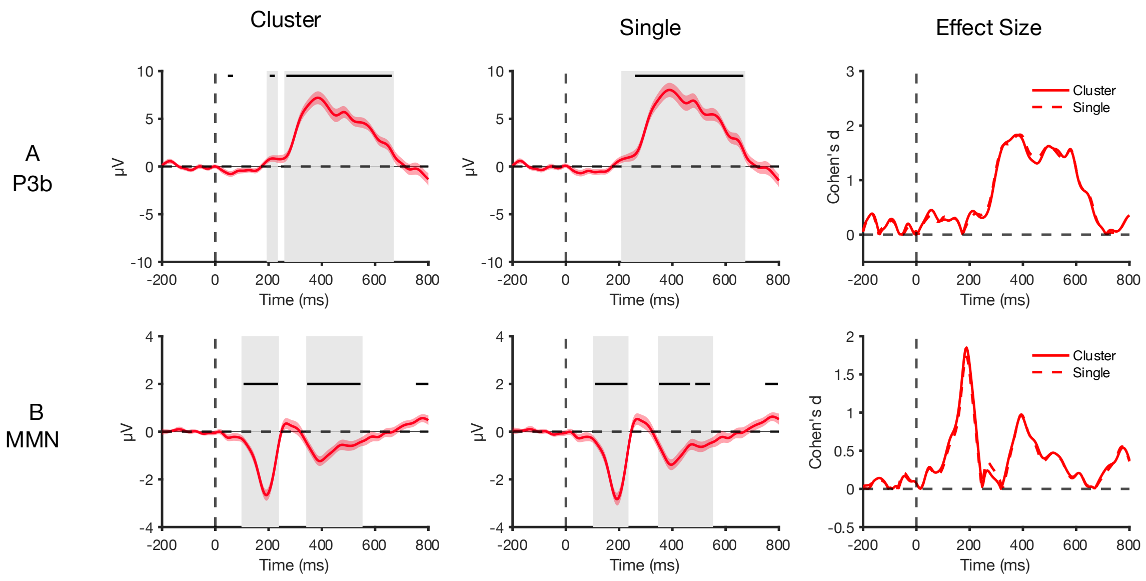


Figure S4. Point-by-point ERP difference voltages, for the components elicited by the oddball paradigms (P3b/MMN) using single sites and clusters of sites. The shaded bands around the waveform show the standard error of the mean. Gray shading indicates significance at $p < 0.05$, FDR corrected. The right column shows the effect size (Cohen's $d_z$) at each time point for each analysis method. The cluster sites tend to yield roughly the same effect sizes as their single site counterparts.

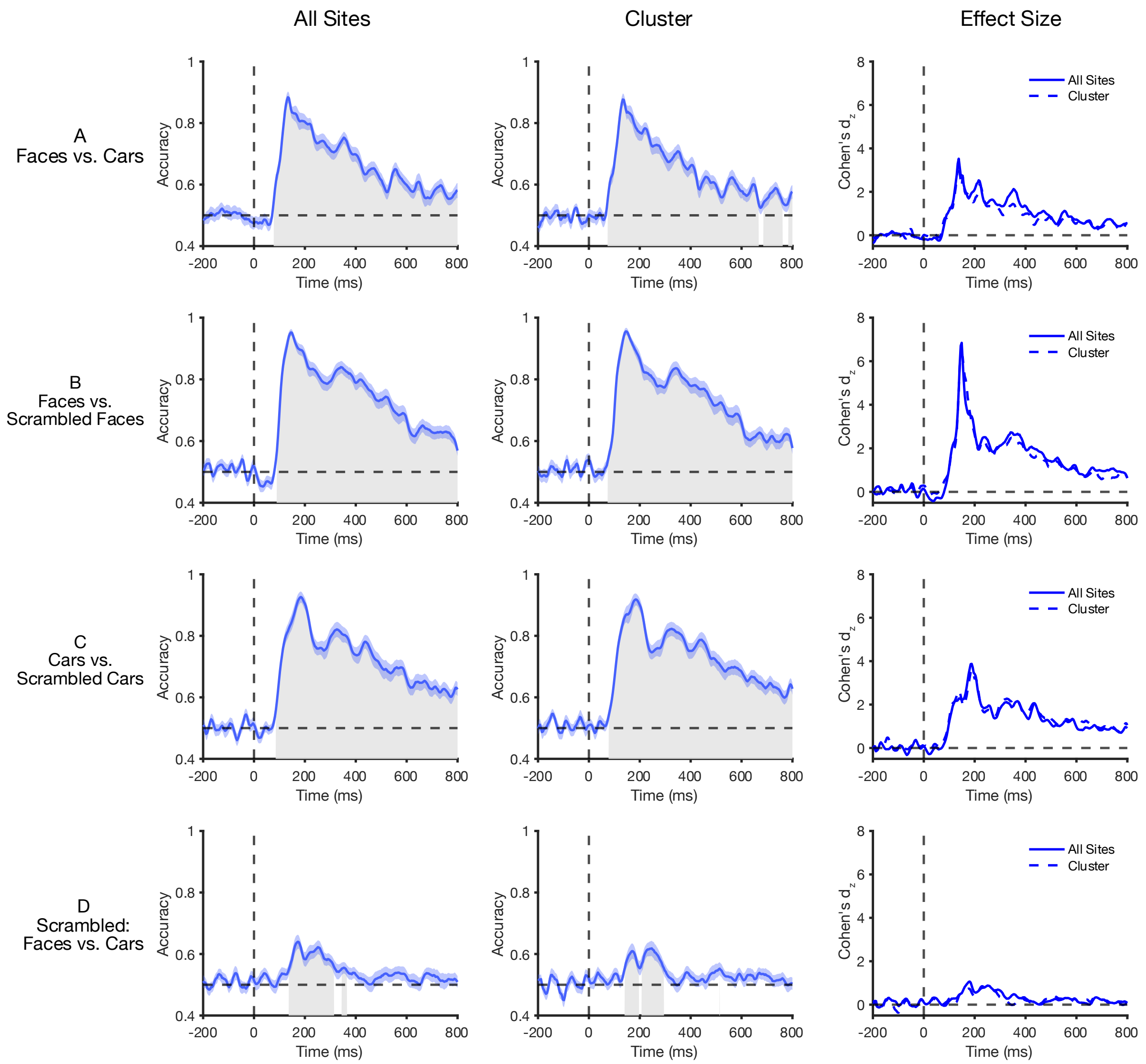


Figure S5. Point-by-point decoding accuracy for the 4 comparisons analogous to the components elicited by the N170 paradigm using clusters of sites, and all sites. The shaded bands around the waveform show the standard error of the mean. Gray shading indicates significance at $p < 0.05$, FDR corrected. The right column shows the effect size (Cohen's $d_z$) at each time point for each analysis method. Using all sites tends to yield larger effect sizes than using a cluster.

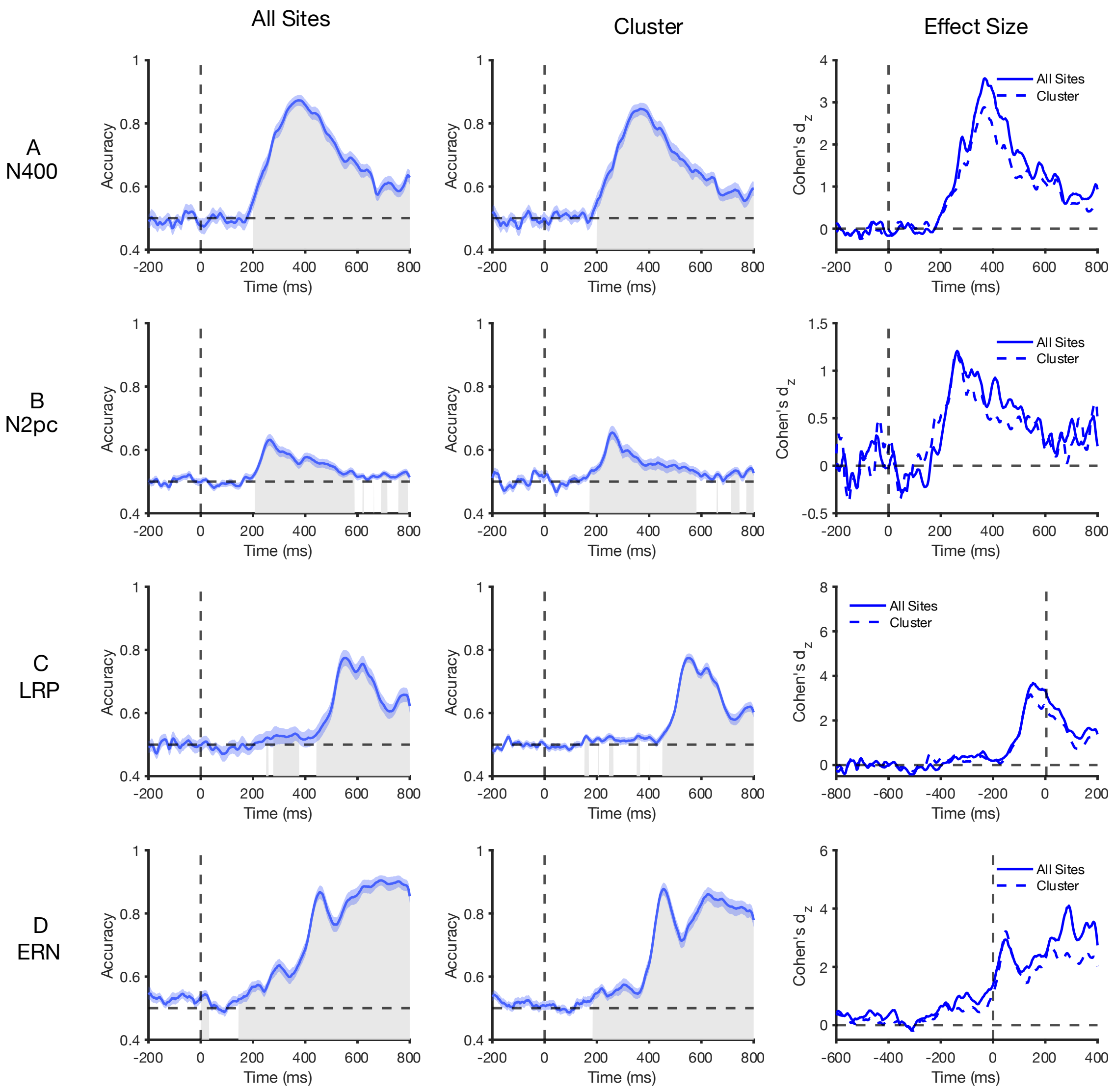


<u>Figure S6.</u> Point-by-point decoding accuracy for the 4 comparisons analogous to the components elicited by the N400, N2pc, LRP/ERN paradigms using clusters of sites, and all sites. The shaded bands around the waveform show the standard error of the mean. Gray shading indicates significance at $p < 0.05$, FDR corrected. The right column shows the effect size (Cohen's $d_z$) at each time point for each analysis method. Using all sites tends to yield larger effect sizes than using a cluster.

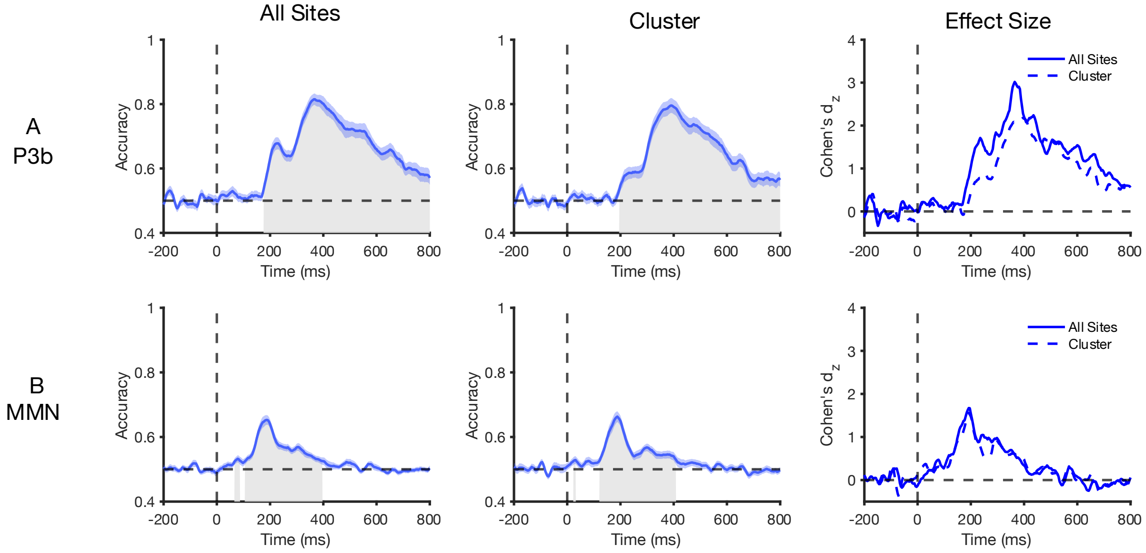


<u>Figure S7.</u> Point-by-point decoding accuracy for the comparisons analogous to the components elicited by the P3b and MMN paradigms using clusters of sites, and all sites. The shaded bands around the waveform show the standard error of the mean. Gray shading indicates significance at $p < 0.05$, FDR corrected. The right column shows the effect size (Cohen's $d_z$) at each time point for each analysis method. Using all sites tends to yield roughly the same effect sizes as using single sites.

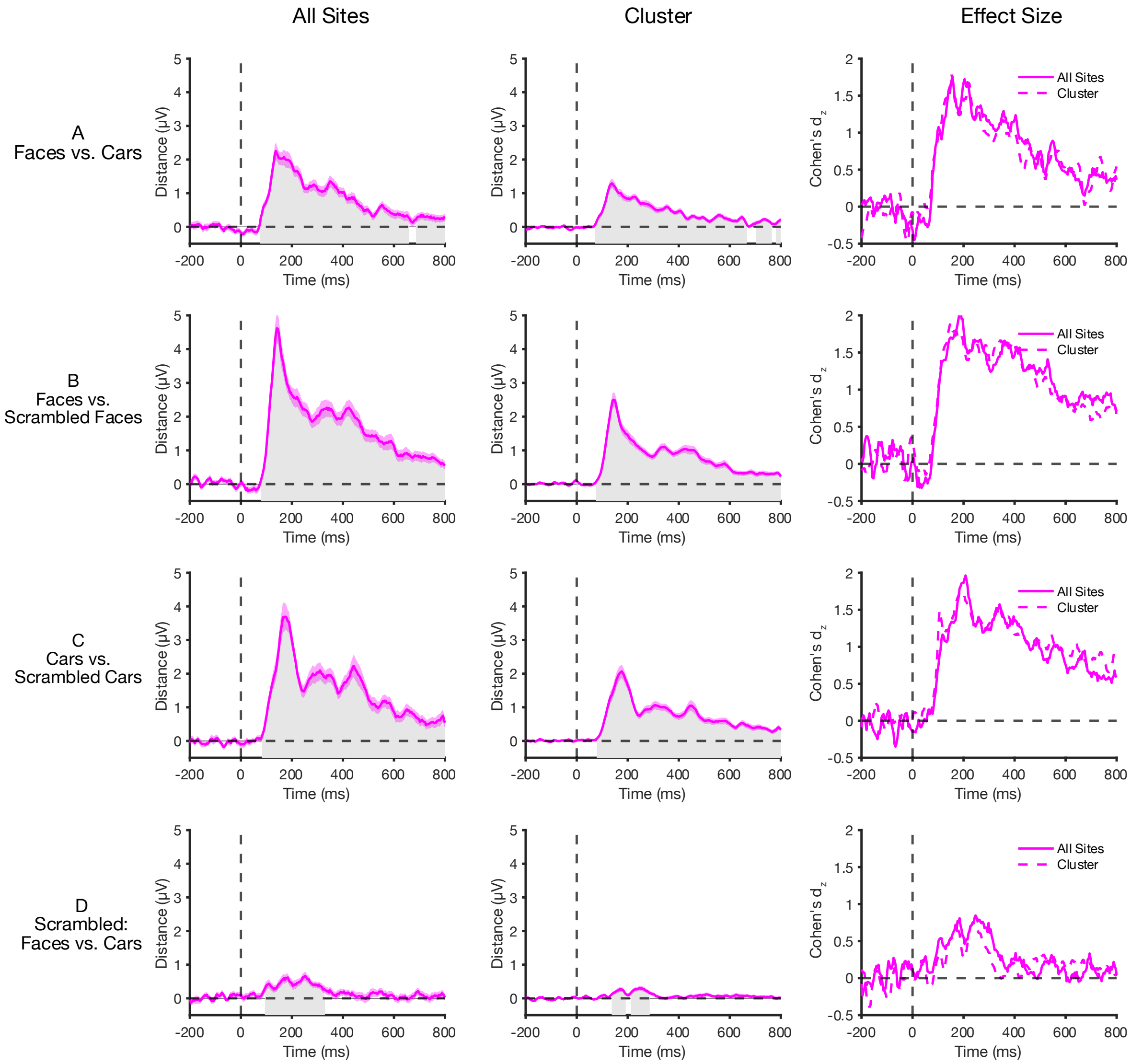


Figure S8. Point-by-point crossnobis distance for the 4 comparisons analogous to the components elicited by the N170 paradigm using clusters of sites, and all sites. The shaded bands around the waveform show the standard error of the mean. Gray shading indicates significance at $p < 0.05$, FDR corrected. The right column shows the effect size (Cohen's $d_z$) at each time point for each analysis method. Using all sites or clusters tend to yield roughly the same effect sizes across the time series.

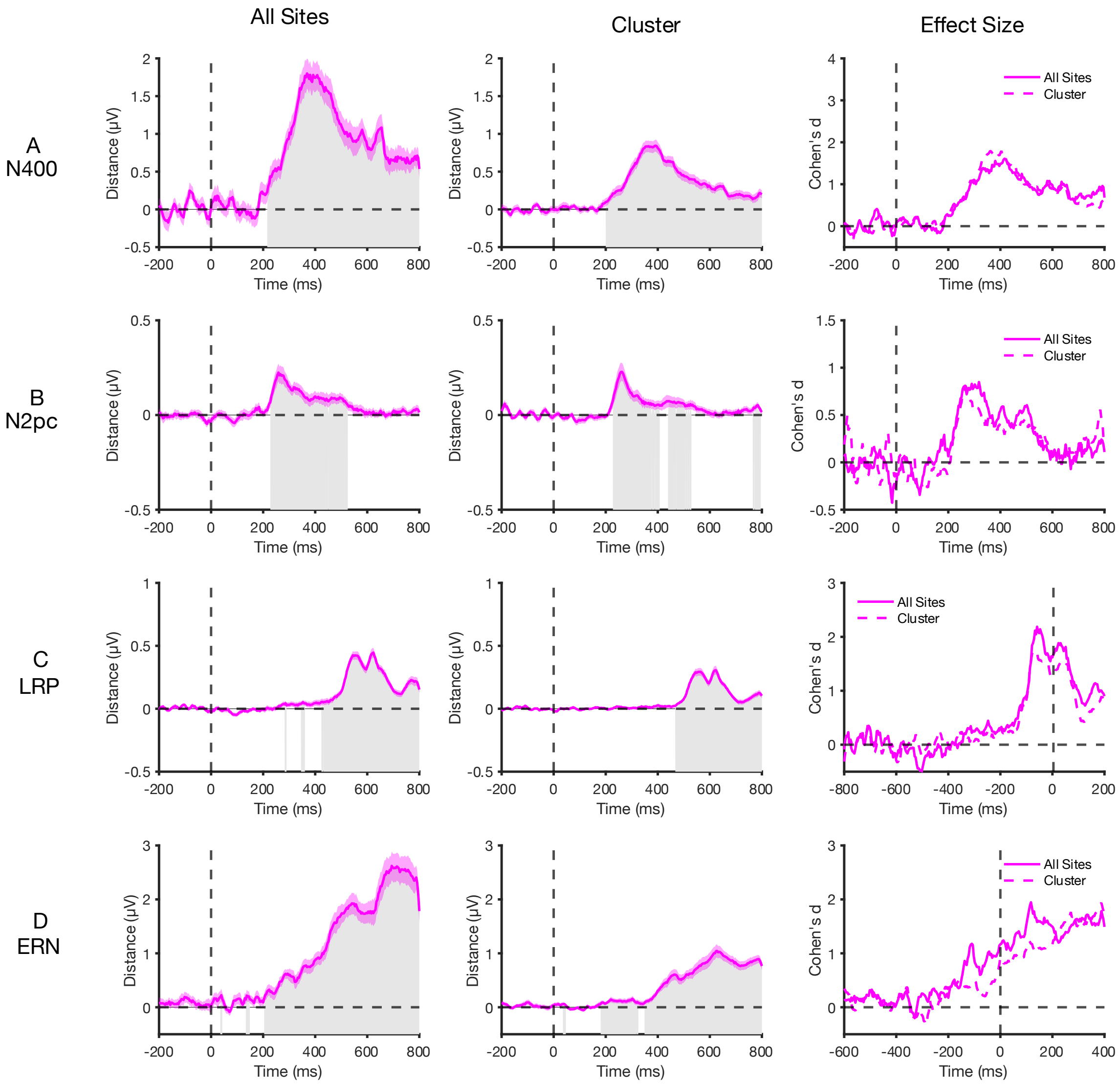


Figure S9. Point-by-point crossnobis distance for the 4 comparisons analogous to the components elicited by the N400, N2pc, LRP/ERN paradigms using clusters of sites, and all sites. The shaded bands around the waveform show the standard error of the mean. Gray shading indicates significance at $p < 0.05$, FDR corrected. The right column shows the effect size (Cohen's $d_z$) at each time point for each analysis method. Using all sites or clusters tend to yield roughly the same effect sizes across the time series.

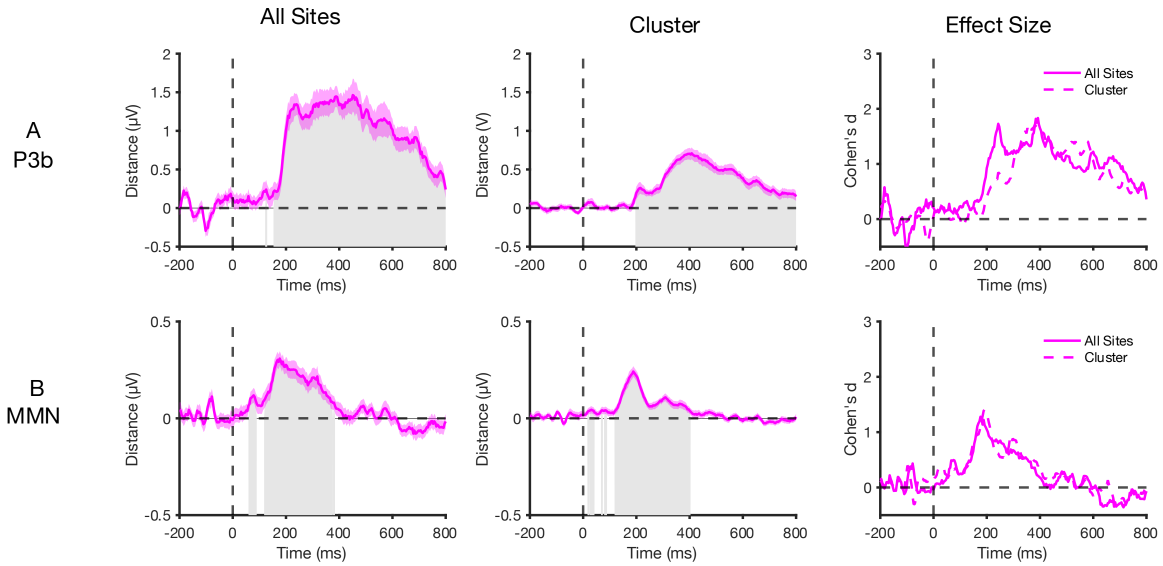


Figure S10. Point-by-point crossnobis distance for the comparisons analogous to the components elicited by the P3b and MMN paradigms using clusters of sites, and all sites. The shaded bands around the waveform show the standard error of the mean. Gray shading indicates significance at $p < 0.05$, FDR corrected. The right column shows the effect size (Cohen's $d_z$) at each time point for each analysis method. Using all sites tends to yield roughly the same effect sizes as using single sites.

Table S1. Summary of univariate analyses when the data were measured from a single channel instead of a cluster of channels.

| | | Univariate Parameters | | | | | |
|---|---|---|---|---|---|---|---|
| ERP Component | Measurement Window | Electrode Measured | *N* | *df* | Mean Amp *t* | Mean Amp *p* | Time Locking Point |
| N170 FvC | 110 to 150 | PO8 | 37 | 36 | 7.36 | p < 0.0001 | stimulus onset |
| N170 FvSF | 110 to 150 | PO8 | 37 | 36 | 7.90 | p < 0.0001 | stimulus onset |
| N170 CvSC | 110 to 150 | PO8 | 37 | 36 | 4.38 | p < 0.0001 | stimulus onset |
| N170 SFvSC | 110 to 150 | PO8 | 37 | 36 | 3.13 | p = 0.004 | stimulus onset |
| N400 | 300 to 500 | CPz | 37 | 36 | 13.96 | p < 0.0001 | stimulus onset |
| N2pc | 200 to 275 | PO7/PO8 | 35 | 34 | 5.86 | p < 0.0001 | stimulus onset |
| LRP | -100 to 0 | C3/C4 | 37 | 36 | 15.56 | p < 0.0001 | response onset |
| ERN | 0 to 100 | FCz | 36 | 35 | 9.44 | p < 0.0001 | response onset |
| P3 | 300 to 600 | Pz | 34 | 33 | 10.82 | p < 0.0001 | stimulus onset |
| MMN | 125 to 225 | FCz | 39 | 38 | 9.49 | p < 0.0001 | stimulus onset |

Table S2. Summary of SVM results when the decoding was performed only on the data from the electrodes used in the main univariate analyses.

SVM Parameters

| ERP Component | Measurement Window | Electrode Sites | Avg # of Decoding Trials | Std Deviation | # of Crossfolds | Min # of Trials Per Crossfold | *N* | *df* | Decoding *t* | Decoding *p* | Time Locking Point |
|---|---|---|---|---|---|---|---|---|---|---|---|
| N170 FvC | 110 to 150 | P7, P8, PO7, PO8, O1,O2 | 68.41 | 6.95 | 5 | 13 | 37 | 36 | 41.00 | $p < 0.0001$ | stimulus onset |
| N170 FvSF | 110 to 150 | P7, P8, PO7, PO8, O1,O2 | 67.43 | 6.87 | 5 | 13 | 37 | 36 | 46.81 | $p < 0.0001$ | stimulus onset |
| N170 CvSC | 110 to 150 | P7, P8, PO7, PO8, O1,O2 | 67.16 | 8.05 | 5 | 13 | 37 | 36 | 40.36 | $p < 0.0001$ | stimulus onset |
| N170 SFvSC | 110 to 150 | P7, P8, PO7, PO8, O1,O2 | 66.65 | 7.63 | 5 | 13 | 37 | 36 | 32.58 | $p < 0.0001$ | stimulus onset |
| N400 | 300 to 500 | Cz, Pz, C3, C4, P3, P4 | 51.05 | 5.08 | 4 | 10 | 37 | 36 | 38.90 | $p < 0.0001$ | stimulus onset |
| N2pc | 200 to 275 | P7/8, PO7/8, PO3/4, O1/2, P3/4 | 57.54 | 9.87 | 5 | 11 | 35 | 34 | 6.68 | $p = 0.0002$ | stimulus onset |
| LRP | -100 to 0 | F3/4, FC3/4, C3/4, C5/6, P3/4 | 84.15 | 7.41 | 5 | 16 | 37 | 36 | 19.42 | $p < 0.0001$ | response onset |
| ERN | 0 to 100 | Fz, FCz, Cz, FC3, FC4 | 41.17 | 21.12 | 4 | 10 | 36 | 35 | 41.82 | $p < 0.0001$ | response onset |
| P3 | 300 to 600 | Cz, CPz, Pz, P3, P4 | 33.62 | 4.05 | 3 | 11 | 34 | 33 | 44.51 | $p < 0.0001$ | stimulus onset |
| MMN | 125 to 225 | Fz, FCz, Cz, FC3, FC4 | 185.77 | 13.94 | 10 | 18 | 39 | 38 | 49.95 | $p < 0.0001$ | stimulus onset |

Supplementary Table 3. Summary of crossnobis results when the decoding was performed only on the data from the electrodes used in the main univariate analyses.

<u>Crossnobis Parameters</u>

| ERP Component | Measurement Window | Electrode Sites | *N* | *df* | Xnobis *t* | Xnobis *p* | Time Locking Point |
|---|---|---|---|---|---|---|---|
| N170 FvC | 110 to 150 | P7, P8, PO7, PO8, O1,O2 | 37 | 36 | 10.59 | $p < 0.0001$ | stimulus onset |
| N170 FvSF | 110 to 150 | P7, P8, PO7, PO8, O1,O2 | 37 | 36 | 11.60 | $p < 0.0001$ | stimulus onset |
| N170 CvSC | 110 to 150 | P7, P8, PO7, PO8, O1,O2 | 37 | 36 | 8.37 | $p < 0.0001$ | stimulus onset |
| N170 SFvSC | 110 to 150 | P7, P8, PO7, PO8, O1,O2 | 37 | 36 | 2.47 | $p = 0.008$ | stimulus onset |
| N400 | 300 to 500 | Cz, Pz, C3, C4, P3, P4 | 37 | 36 | 11.19 | $p < 0.0001$ | stimulus onset |
| N2pc | 200 to 275 | P7/8, PO7/8, PO3/4, O1/2, P3/4 | 35 | 34 | 4.14 | $p < 0.0001$ | stimulus onset |
| LRP | -100 to 0 | F3/4, FC3/4, C3/4, C5/6, P3/4 | 37 | 36 | 11.14 | $p < 0.0001$ | response onset |
| ERN | 0 to 100 | Fz, FCz, Cz, FC3, FC4 | 36 | 35 | 6.58 | $p < 0.0001$ | response onset |
| P3 | 300 to 600 | Cz, CPz, Pz, P3, P4 | 34 | 33 | 11.33 | $p < 0.0001$ | stimulus onset |
| MMN | 125 to 225 | Fz, FCz, Cz, FC3, FC4 | 39 | 38 | 8.58 | $p < 0.0001$ | stimulus onset |
